# High prevalence of multi-drug resistant bacteria in faecal samples from UK passerine birds

**DOI:** 10.1101/2024.07.24.604896

**Authors:** Jenny C. Dunn, Simon R. Clegg

## Abstract

Wild birds are a near ubiquitous sight in gardens, offering pleasure to many people through supplementary feeding, song, or other interactions. However, they are also potential carriers of many pathogens, including *Campylobacter*, *Salmonella, Enterococcus* and *E. coli*; some of these may be resistant to commonly used drugs. This study collected faecal samples from multiple species of UK passerine birds, isolating bacterial pathogens to assess carriage and drug resistances associated with those bacteria. 75% of birds were carrying at least one bacterial species which was multi drug resistant (MDR; resistant to three or more classes of antimicrobial), with 11.6% of birds carrying *Salmonella* spp., 18.9% carrying *Campylobacter* spp., 78% carrying *Enterococcus* spp., and all carrying *E. coli* strains. Many of these strains were shown to be MDR with 70%, 88%, 32% and 59% respectively. Intercontinental migration was shown to be a risk factor for carriage of many of the pathogens, as was an associated with human habitation. Age was also a risk factor with younger birds twice as likely to carry *Campylobacter*spp. than adults, and house sparrows (*Passer domesticus*) and blackbirds (*Turdus merula*) being particularly high-level carriers compared to other species. The high-level carriage and shedding of MDR *E. coli* and other zoonotic pathogens within the faecal samples of multiple species of passerine birds offers a timely reminder of the risks which these bacteria, and their drug resistance profiles may pose to human and animal health in the UK and worldwide. It also shows a level of high environmental contamination, which birds may continue to contribute towards, until our use of antibiotics, and level of drug resistant bacteria is decreased. Developing mechanisms for reducing levels of carriage of MDR bacteria in wild bird populations through, for example, increased hygiene around bird feeding practices, may be key in reducing environmental contamination.

**Highlights:**

- 75% of wild birds were carrying at least one MDR bacterium
- Young birds were twice as likely to carry *Campylobacter* than adults
- House sparrows and blackbirds were particularly likely to carry *Campylobacter*
- All intercontinental migrants with *Salmonella* carried MDR strains

## Introduction

Antimicrobial resistant (AMR) pathogens are of increasing concern worldwide within both the veterinary and human medical fields making it a One Health concern (Robinson et al., 2016). Often caused by our overuse and misuse of antimicrobials, AMR bacteria are found in almost every ecosystem investigated, from animals and humans through to soil and water (reviewed by Velazquez-Mesa et al (Velazquez-Meza et al., 2022)). This in turn has an effect of increasing both morbidity and mortality in domesticated animals and humans, as well as increases in costs for treatments (Edwards et al., 2019). Among the most under-investigated systems for AMR carriage is wildlife.

Wildlife pose a major risk for carriage and transmission of AMR bacteria due to their indiscriminate defaecation, and potential to cover large distances, potentially shedding AMR bacteria across wide areas leading to environmental contamination (Arnold et al., 2016). As treatment of wildlife with any antimicrobial is uncommon, these species act as a good indicator of the levels of contamination of the environment with AMR, with wild animals and birds encountering AMR bacteria through food or water (Swift et al., 2019). Worldwide, wild birds offer a large amount of pleasure to people, with as many as 75% of households encouraging them into their gardens with supplementary feeding stations (Robb et al., 2008). It has been suggested that AMR in wild birds and other wildlife is associated with anthropogenic activities and environments (Swift et al., 2019). However, wild birds are widely considered as potential disseminators of AMR bacteria, due to their tendency to migrate long distances, and to occupy a range of habitats known to be contaminated (Elsohaby et al., 2021). Passerine birds within the UK inhabit all available ecosystems, survive on different diets, and have different migration patterns, so they may have a high risk of introduction of novel pathogens into the UK, and subsequent widescale dissemination.

*Escherichia coli* is one of the most commonly tested bacterial species for AMR carriage, due to its simple and rapid isolation, as well as its prevalence as a major component of the gut microbiota in many animal species (Anjum et al., 2021). In addition, this bacterial species also has a high tendency to both acquire and lose antimicrobial resistance genes. Birds are well known carriers of several different pathogens, including *Campylobacter* spp. and *Salmonella* spp., both of which are major concern for both human and veterinary medicine, causing a range of symptoms from gastrointestinal disease to abortion depending on the specific bacterial species (Alley et al., 2002; Cody et al., 2015). *Enterococcus* spp. is also a major issue within human medicine, being associated with urinary tract infections, septicaemia, and infected wounds (Vu and Carvalho, 2011). Given the potential severity of these pathogens, the presence of AMR poses an increased risk to animal and human health. Indeed, bacterial pathogens of wild avian origin have been suggested to be the cause of outbreaks of disease in humans (Alley et al., 2002; Cody et al., 2015). Many human *Campylobacter* spp., *Salmonella* spp., and *Enterococcus* isolates associated with disease also show some level of AMR. Therefore, studies on bacterial carriage, prevalence and antimicrobial susceptibility are important to inform optimal treatment regimes for human and animal diseases.

Many studies of AMR bacterial carriage in wild birds focus on waterfowl, as these are major risk factors for the transmission of avian influenza (McDuie et al., 2022). Previous studies have shown AMR *E. coli* in birds from many different countries, as well as the carriage of *Salmonella* spp., *Campylobacter* spp. and *Enterococcus* spp. However, despite their near ubiquitous presence, little is known about the bacterial pathogen carriage of songbirds (Passeriformes) within the UK, which is a major site for migration for many birds across the world (Sparks et al., 2007), bringing with it the potential risk of introduction of new or novel pathogens, and or AMR genes. In this study, we screen 259 faecal samples from 23 species of passerine birds to quantify the prevalence of four different bacterial pathogens and assess their susceptibility to a range of different antimicrobials from different classes which are used to treat both animal and human clinical cases. We then test for host and ecological associations with infection by each pathogen to elicit potential drivers or risk factors for infection.

## Methods

### Sites and sample collection

Faecal sample were collected from wild birds caught as part of standard bird ringing activities at three different sites. One site, near Braintree, Essex, UK (51°53’24.8”N, 0°33’18.3”E) was a residential garden of approximately 0.75 ha surrounded by arable farmland, where ten birdfeeders were provided to encourage birds into the garden. Feeders were kept full year-round, with provided food including sunflower hearts (*Helianthus annuus*), peanuts (*Arachis hypogaea*), a bird seed mix (dominated by wheat (*Triticum aestivum*), and nyjer seed (*Guizotia abyssinica*). The second site, near Potterhanworth, Lincolnshire, UK (53°11’02.5”N, 0°25’21.5”W) was a small woodland copse surrounded by arable farmland, with wheat provided year-round to feed gamebirds (mostly ring-necked pheasants (*Phasianus colchicus*)). The third site, near Glentham, Lincolnshire, UK (53°24’03.8”N, 0°29’37.5”W) consisted of three small lakes bordered by scrub (mostly hawthorn (*Crataegus* sp.) and blackthorn (*Prunus spinosa*) surrounded by arable farmland, where no supplementary food was provided.

At each site, birds were captured using mist nets on days that were dry and still. Birds were caught on 14 occasions per site between June - August 2022 and fitted with an individually numbered BTO metal ring before being aged and sexed where possible according to plumage characteristics (Svensson, 1992), measured (maximum wing chord measured using a slotted wing rule, ± 0.5mm) and weighed using a digital balance (± 0.1 g). Faecal samples were collected following release of the bird, from the inside of a clean and disinfected cotton bird bag, stored at ambient temperature in the field (up to 6 hours) and then stored at 4°C until processing.

### *E. coli* isolation

From each faecal sample, 0.1g was resuspended in 900µl of sterile physiological saline (Melford, UK) before being plated onto MacConkey agar (Oxoid, UK) and incubated for 24 hours at 37°C. Three suspected *E. coli* colonies were selected from each plate and resuspended in PBS before being subcultured onto Columbia agar plates (Oxoid, UK) and incubated for 24 hours at 37°C to obtain a monoculture. Bacterial strains were identified using Gram staining and PCR to confirm that isolates were *E. coli* (Sabat et al., 2000). A selection of these were subjected for sequencing to confirm PCR specificity (data not shown). All strains were stored with nutrient broth (Oxoid, UK) and glycerol (Sigma Aldrich UK) at a ratio of 80:20 at -80°C until further analysis.

Antimicrobial susceptibility was performed using the Kirby-Bauer disk diffusion method according to the European Committee on Antimicrobial Susceptibility Testing (EUCAST) guidelines (Matuschek et al., 2014). Isolates were recovered from the frozen stocks and spread onto Columbia Agar with 5% sheep blood (Scientific Laboratory Supplies, UK) before being suspended in 0.8% saline solution to obtain a turbidity of 0.5 McFarland units. This inoculum was transferred onto Mueller-Hinton Agar (Oxoid, UK) and antimicrobial discs placed on the surface. In total, 22 antimicrobials from 11 different classes were tested based on previous studies (Guenther et al., 2010; Ong et al., 2020) to allow for comparison. Plates were incubated for 18-20 hours at 36°C before sensitivity or resistance was assessed through growth inhibition diameter according to EUCAST breakpoints (EUCAST). Exceptions to this were ceftiofur, enrofloxacin and tetracycline which were evaluated based on previous studies (Magiorakos et al., 2012; Markey et al., 2013). Multidrug resistance (MDR) was determined if an isolate was shown as fully resistant to at least one antimicrobial agent in three or more antimicrobial classes (Magiorakos et al., 2012). *E. coli* ATCC 11755 was used as a reference strain.

### Salmonella isolation

Samples were analysed for *Salmonella* spp., based on ISO 6579-1:2017. Firstly, 0.1g of each faecal sample was pre-enriched in 1:10 buffered peptone water (ThermoFisher Scientific, UK) before incubation at 37°C for 18-20 hours. From this pre-enrichment, 100µl were transferred onto semi solid modified Rappaport Vassiliadis (Difco, UK) before incubation for 48 hours at 41.5°C. Colonies suggestive of *Salmonella* spp., were further inoculated onto Xylose-Lysine Deoxycholate (ThermoFisher Scientific, UK) and chromogenic agar specific for detection of C8-esterase activity (ASAP, bioMerieux, Marcy l’Étoile, France) and incubated at 37°C for 48 hours. Isolates were confirmed to be *Salmonella* spp. by PCR (DePaola et al., 2010), and serotyped using the antigenic agglutination method with specific antisera according to the White-Kauffmann-Le Minor scheme (ISO 6579-1:2017).

### Antimicrobial resistance testing

Antimicrobial resistance was determined using ISO 20776-1:2006 using a Mueller Hinton broth microdilution (Oxoid, UK) to allow for determination of the minimum inhibitory concentration (MIC) using EUCAST breakpoints (Grimont and Weill, 2007; Leclercq et al., 2013). Isolates were tested against ten different antimicrobials from seven different classes (amikacin, amoxicillin-clavulanic acid, ampicillin, ceftriaxone, chloramphenicol, gentamycin, nalidixic acid, norfloxacin, sulfamethoxazole-trimethoprim and tetracycline). After incubation of 24 hours at 37°C the inhibition zone around each disc was measured and interpreted using EUCAST and CLSI breakpoints. A multidrug-resistant (MDR) strain was considered when the isolate was non-susceptible to at least one antimicrobial agent in three or more antimicrobial categories (Schwarz et al., 2010).

### Enterococcus isolation

Faecal dilutions were inoculated onto Slanetz and Bartley agar (Oxoid, UK) and incubated at 37°C for 48 hours. A single isolate (red, maroon or pink coloured colony) was removed from the plate and subcultured in LB broth (Melford, UK) before incubation at 37°C for 48 hours. DNA from a small aliquot of this culture was extracted using a boil preparation (Peng et al., 2013) and was confirmed to be *Enterococcus* spp. using PCR (Ke et al., 1999). Speciation was carried out using the methods described by Jackson et al., (2004).

### Antimicrobial susceptibility testing

Antimicrobial minimum inhibitory concentrations were calculated for all *Enterococcus* isolates following the CLSI guidelines. Susceptibilities were determined to vancomycin, teicoplanin, ampicillin, streptomycin, gentamycin, kanamycin, chloramphenicol, tetracycline, erythromycin and ciprofloxacin in line with previous studies (Santos et al., 2013; Cagnoli et al., 2022) to allow for comparisons. For controls, the reference strains used were *E. faecalis* ATCC 29212 and *S. aureus* ATCC 29213. Analysis compared the breakpoints to those with the CLSI standards or the national antimicrobial resistance monitoring system.

### Campylobacter isolation

Isolation of *Campylobacter* spp., was performed using the ISO 10272-1:2017 as described previously (Mencía-Gutiérrez et al., 2021). Briefly, modified charcoal cefoperazone deoxycholate (mCCDA) and Preston agar (Oxoid, UK) were both streaked with the samples, and incubated in a microaerophilic environment at 41.5°C for 48 hours. Colonies indicative of *Campylobacter* spp., were examined by PCR for genus, and species confirmation (Wang et al., 2002).

### AMR testing

All *Campylobacter*isolates were subjected to antimicrobial resistance testing using the broth microdilution method (Luber et al., 2003). Using sensitive *Campylobacter* EUCAMP2® plates (ThermoFisher Scientific, UK) according to manufacturers instructions, each isolate was tested against 10 different antimicrobials : azithromycin, chloramphenicol, ciprofloxacin, enrofloxacin, erythromycin, gentamycin, nalidixic acid, tetracycline, trimethoprim-sulfamethoxazole and amoxicillin. Susceptibility was determined based on the epidemiological cut off values established by EUCAST.

### Statistical analyses

All statistical analyses were conducted in R version 4.3.1 “Beagle Scouts” for Mac (R Core Team, 2021). Two binomial general linear models were constructed to test for associations between each of environmental variables and host ecological variables on the presence and MDR status of each bacterium (fourteen models in total; all birds were carrying *E. coli* so no models were constructed to test for associations with *E. coli* presence). For each model, the binomial response variable was the presence or absence of either the bacterium, or MDR. For the environmental model, fixed factors comprised Site (a 3-level factor), Species (a 16-level factor), Age (a two-level factor of juvenile (hatched during the calendar year of capture) or adult (hatched prior to this) and day (a continuous variable). For the ecological models, fixed factors comprised Migrant status (resident or long-distance migrant; species that may undertake short-distance migration were classified as resident for the purposes of this analysis), whether the species was associated with human habitation (a two-level factor of Yes or No), whether the species was granivorous (Yes or No) or insectivorous (Yes or No) and the number of food types used by the species (a continuous variable). Host ecological data were extracted from (Storchová and Hořák, 2018) at the species level. Models were simplified by removing the least significant term in turn (as determined by likelihood ratio tests) until either all remaining terms in the model were significant at p<0.1, or only the null model remained.

## Results

### Salmonella

*Salmonella* spp. was identified from 30 (11.6%) of 259 faecal samples using PCR. Twelve serovars of *Salmonella* were isolated (Table 1), the most common of which, *Salmonella enterica*serovar Typhimurium, was isolated from eleven individuals from seven bird species (Table 1). Multi-drug resistance (MDR), defined as resistance to at least three classes of antibiotic, was identified in 21 (70%) of the 30 positive samples (Table 1). MDR was identified in the serovars Agona (100%, n=2), Havana (100%, n=1), Meleagridis (100%, n=2), Muenchen (50%, n=1), Muenster (100%, n=2), Newport (50%, n=2), Panama (50%, n=2), Parathyphi B var. Java (33%, n=3), Rissen (100%, n=1) and Typhimurium (82%, n=11; Table 1).

**Table 1.** The number of individuals from each host species from which each Salmonella strain was isolated. The number of individuals with strains showing MDR is provided in parentheses.

| *Salmonella* strain | Host species |  |  |  |  |  |  |  |  |  |  |
| --- | --- | --- | --- | --- | --- | --- | --- | --- | --- | --- | --- |
|  | Blackbird | Blackcap | Blue tit | Chaffinch | Chiffchaff | Dunnock | Goldfinch | House sparrow | Robin | Whitethroat | Wren |
| Agona |  |  |  |  |  |  |  | 2 (2) |  |  |  |
| Havana |  |  |  |  |  |  |  |  |  |  | 1 (1) |
| Heidelberg | 1 (0) |  |  |  |  |  |  |  |  |  |  |
| Meleagris |  |  |  |  |  | 1 (1) |  | 1 (1) |  |  |  |
| Montevideo |  |  |  |  |  |  |  |  | 1 (0) |  |  |
| Muenchen |  | 1 (1) | 1 (0) |  |  |  |  |  |  |  |  |
| Muenster |  | 1 (1) | 1 (1) |  |  |  |  |  |  |  |  |
| Newport | 1 (0) | 1 (1) |  |  |  |  |  |  |  |  |  |
| Panama |  |  |  |  |  |  |  | 2 (1) |  |  |  |
| Paratyphi B var. |  |  |  |  |  | 1 (0) | 1 (1) |  | 1 (0) |  |  |
| Java |  |  |  |  |  |  |  |  |  |  |  |
| Rissen |  |  |  |  |  |  | 1 (1) |  |  |  |  |
| Typhimurium |  | 2 (2) | 1 (1) | 2 (1) | 1 (1) |  |  | 3 (2) |  | 1 (1) | 1 (1) |

Resistance was highest to tetracycline (n=21, 70%), chloramphenicol (n=17, 57%) and ampicillin (n=15, 50%), with 50% or more of samples showing complete resistance (Table S1). Resistance was lowest to nalidixic acid, with no samples showing complete resistance and three samples (10%) showing intermediate levels of resistance, followed by amikacin, where 1 sample (3%) showed complete resistance and no samples showed intermediate resistance (Table S1).

Neither the presence of *Salmonella*, nor the presence of MDR *Salmonella* within positive samples, differed between sites, or between adult and juvenile birds, and did not vary with Julian day (Table S2). The presence of *Salmonella* did not differ between species (Table S2a; this analysis was not conducted for MDR *Salmonella* due to small sample sizes), and none of diet, migration strategy or breeding presence within human settlements influenced the presence of *Salmonella* (Table 2). However, all intercontinental migrants with *Salmonella* were carrying strains with MDR (Migrants: 100% MDR [n=7], non-migrants: 59% MDR [n=22]; Table 2), and birds carrying non-MDR *Salmonella* had higher diet diversity than those carrying MDR *Salmonella* (non-MDR *Salmonella*: 2.22 ± 0.15 food types; MDR *Salmonella*: 1.75 ± 0.10 food types; Table 2).

**Table 2.** Results of binomial generalised linear models testing the effect of host species ecological traits on the presence of a) *Salmonella* and b) MDR *Salmonella* in avian faecal samples.

| a) *Salmonella* | Dev | df | p |
| --- | --- | --- | --- |
| Human habitation | 0.496 | 1, 254 | 0.481 |
| Granivorous | 1.162 | 1, 253 | 0.281 |
| Insectivorous | 0.561 | 1, 252 | 0.454 |
| Diet diversity | 0.592 | 1, 251 | 0.442 |
| Migratory (Y/N) | 0.104 | 1, 250 | 0.747 |
| b) MDR *Salmonella* | Dev | df | p |
| **Diet diversity** | **11.353** | **1, 26** | **<0.001** |
| **Migratory** | **9.00** | **1, 26** | **0.003** |
| *Granivorous* | 3.68 | 1, 26 | 0.055 |
| Human habitation | 0.200 | 1, 25 | 0.655 |
| Insectivorous | 0.001 | 1, 24 | 0.999 |

### Campylobacter spp

*Campylobacter* spp. were identified from 49 (18.9%) of 259 faecal samples using PCR (Table 3). Species-specific PCRs identified 3 *C. coli* infections; 22 *C. lari* infections and 24 *C. jejuni* infections; no birds were infected by multiple *Campylobacter* species. Antimicrobial resistance to at least three classes of antibiotic was identified in 43 (88%) of the 49 positive samples (3 (100%), *C. coli* infections; 20 (91%), *C. lari* infections; 20 (83%) and *C. jejuni* infections; Table 3).

**Table 3.** The number of individuals from each host species from which *Campylobacter* spp. was isolated. The number of individuals with strains showing MDR is provided in parentheses. Bullfinch (n=3), dunnock (n=14), garden warbler (n=1), lesser whitethroat (n=1), long-tailed tit (n=3), reed bunting (n=5), sedge warbler (n=3) and yellowhammer (n=4) were tested but found to be negative for *Campylobacter*; these species are not included.

| Host species | *C. coli* | *C. lari* | *C. jejuni* |
| --- | --- | --- | --- |
| Blackbird (n=6) |  | 1 (1) | 2 (2) |
| Blackcap (n=25) |  | 2 (2) | 1 (0) |
| Blue tit (n=29) |  | 4 (4) | 1 (1) |
| Chaffinch (n=30) | 1 (1) | 2 (2) |  |
| Chiffchaff (n=3) |  |  | 1 (1) |
| Goldfinch (n=4) |  | 1 (1) |  |
| Great tit (n=13) |  |  | 1 (1) |
| Greenfinch (n=4) |  |  | 1 (1) |
| House sparrow (n=40) | 1 (1) | 7 (5) | 11 (8) |
| Reed warbler (n=10) |  |  | 1 (1) |
| Robin (n=24) | 1 (1) |  | 3 (3) |
| Song thrush (n=1) |  | 1 (1) |  |
| Whitethroat (n=15) |  | 2 (2) | 1 (1) |
| Willow warbler (n=5) |  |  | 1 (1) |
| Wren (n=16) |  | 2 (2) |  |

Resistance was highest to amoxicillin (n=30, 61%), tetracycline (n=29, 59%) and erythromycin (n=29, 59%), with over 50% of samples showing resistance to all tested antimicrobials (Table S3). Resistance was lowest to trimethoprim-sulfamethoxazole, with 7 samples (14%) showing complete resistance, and a further eight samples (16%) showing intermediate resistance, and to enrofloxacin, where nine samples (18%) showed complete and 8 samples (16%) showed intermediate resistance (Table S3).

The prevalence of Campylobacter differed between species (Table 4a; Figure 1), and juvenile birds were more than twice as likely to be infected as adults (juveniles: 21.5 ± 2.9% prevalence; 8.6 ± 4.8% prevalence). The presence of MDR Campylobacter spp., differed marginally between sites, with 100% (n=9) of positive samples from the fed farmland site being resistant to at least three classes of antibiotic (Table 4). The lowest prevalence of MDR was at the Essex garden site (78% of positive samples; n=23), with 94% (n=17) of positive samples at the unfed farmland site showing MDR.

**Figure 1.**
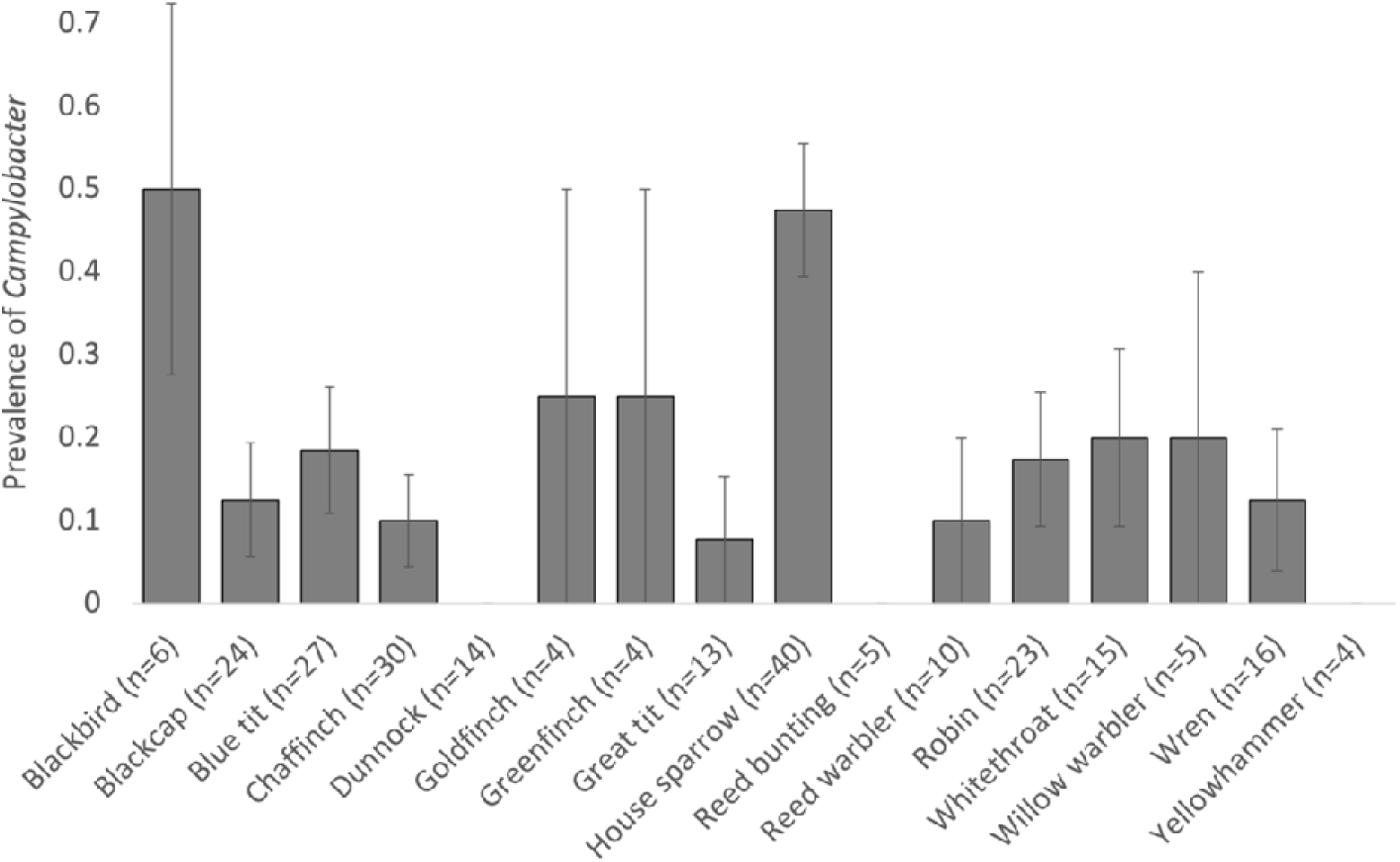
Differences between species in Campylobacter prevalence. Bars show mean ± 1 SE.

**Table 4.**
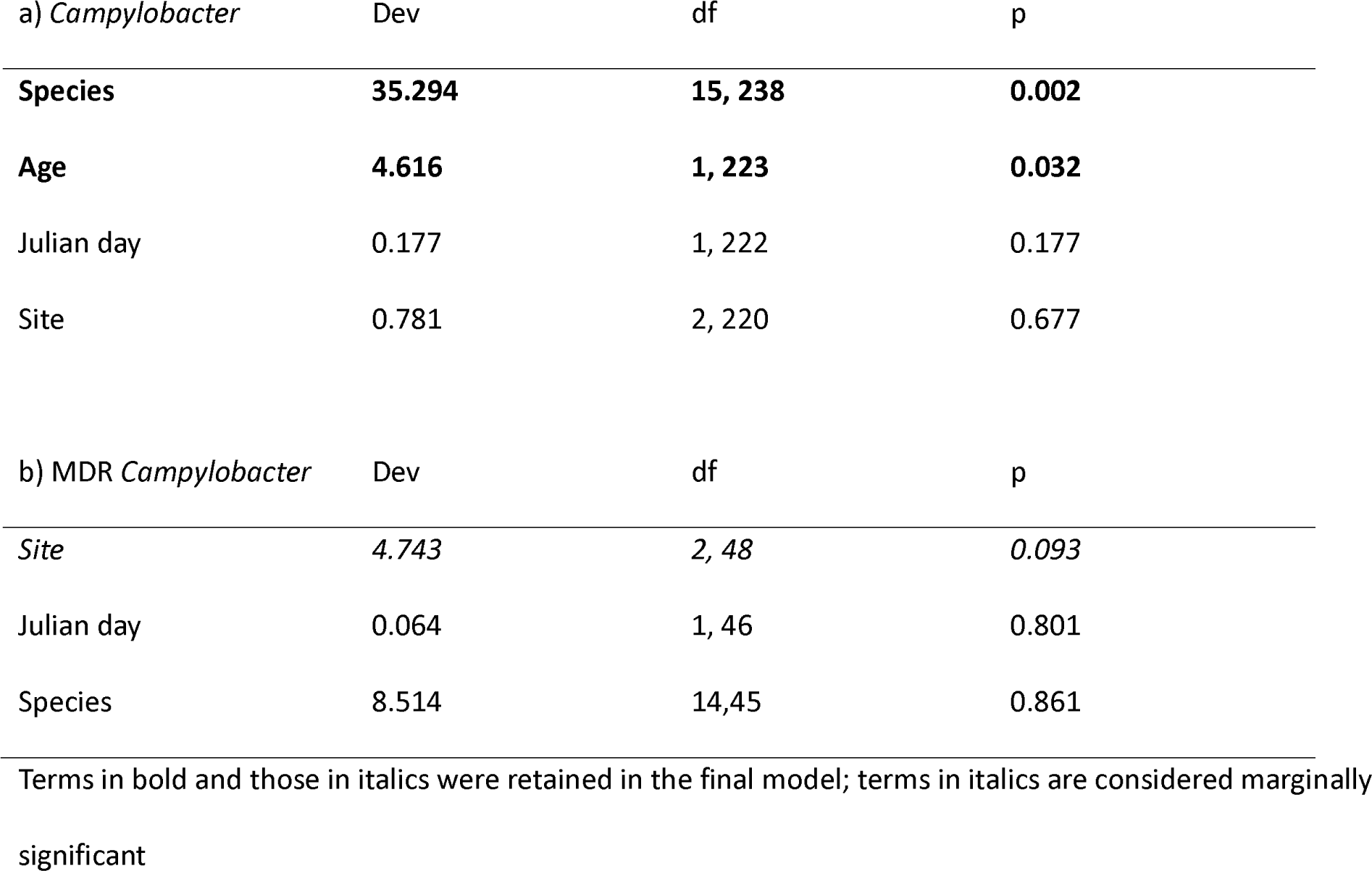
Results of binomial generalised linear models testing the effect of environmental variables on the presence of a) *Campylobacter* and b) MDR *Campylobacter* in avian faecal samples. Age was excluded as a term for b) due to small sample sizes.

| a) *Campylobacter* | Dev | df | p |
| --- | --- | --- | --- |
| **Species** | **35.294** | **15, 238** | **0.002** |
| **Age** | **4.616** | **1, 223** | **0.032** |
| Julian day | 0.177 | 1, 222 | 0.177 |
| Site | 0.781 | 2, 220 | 0.677 |
|  |  |  |  |
| b) MDR *Campylobacter* | Dev | df | p |
| *Site* | *4.743* | *2, 48* | *0.093* |
| Julian day | 0.064 | 1, 46 | 0.801 |
| Species | 8.514 | 14,45 | 0.861 |

Terms in bold and those in italics were retained in the final model; terms in italics are considered marginally significant

The prevalence of *Campylobacter* was higher in birds associated with human habitation (Table 5; associated with human habitation: 69 ± 7 % [n=49]; not associated with human habitation: 44 ± 3 % [n=206]). The prevalence of MDR *Campylobacter* was not associated with any host ecological traits (Table 5).

**Table 5.** Results of binomial generalised linear models testing the effect of host ecological traits on the presence of a) *Campylobacter* and b) MDR *Campylobacter* in avian faecal samples.

| a) *Campylobacter* | Dev | df | p |
| --- | --- | --- | --- |
| **Human habitation** | **13.100** | **1, 253** | **<0.001** |
| *Granivorous* | *3.250* | *1, 253* | *0.072* |
| Diet diversity | 0.051 | 1, 252 | 0.821 |
| Insectivorous | 0.129 | 1, 251 | 0.720 |
| Migratory | 0.015 | 1, 250 | 0.903 |

| b) MDR *Campylobacter* | Dev | df | P |
| --- | --- | --- | --- |
| Human habitation | 0.691 | 1, 48 | 0.406 |
| Migratory | 1.067 | 1, 47 | 0.301 |
| Granivorous | 1.361 | 1, 46 | 0.243 |
| Insectivorous | 0.858 | 1, 45 | 0.384 |
| Diet diversity | 0.872 | 1, 44 | 0.350 |

### Enterococcus spp

*Enterococcus* spp. was identified from 203 (78%) of 259 faecal samples using PCR (Table 6). Species-specific PCRs identified 7 (3.4 % of positives) *E. casseliflavus* infections, 10 (4.9%) *E. durans* infections, 86 (42.4%) *E. faecalis* infections, 88 (43.3%) *E. faecium* infections and 12 (5.9%) *E. hirae* infections. MDR was identified in 65 (32%) of positive infections. No MDR was found in *E. casseliflavus*, *E. durans* or *E. hirae*, but 44 (51.2%) of *E. faecalis*infections and 21 (23.9%) *E. faecium* infections showed MDR (Table 6).

**Table 6.** The number of individuals from each host species from which *Enterococcus* spp. was isolated. The number of individuals with strains showing MDR is provided in parentheses.

|  | *E. casseliflavus* | *E. durans* | *E. faecalis* | *E. faecium* | *E. hirae* |
| --- | --- | --- | --- | --- | --- |
| Blackbird |  |  | 3 (2) |  |  |
| Blackcap | 1 (0) | 2 (0) | 10 (5) | 6 (1) |  |
| Blue tit | 1 (0) | 1 (0) | 13 (7) | 7 (2) | 2 (0) |
| Bullfinch |  | 1 (0) |  | 2 (0) |  |
| Chaffinch | 1 (0) |  | 10 (6) | 11 (1) | 2 (0) |
| Chiffchaff |  |  | 1 (0) |  |  |
| Dunnock |  | 1 (0) | 5 (4) | 6 (2) |  |
| Goldfinch |  |  | 1 (1) | 1 (0) | 1 (0) |
| Great tit |  | 1 (0) | 3 (2) | 3 (0) | 1 (0) |
| Greenfinch |  |  | 1 (0) | 3 (0) |  |
| House sparrow | 2 (0) | 3 (0) | 10 (6) | 13 (9) | 3 (0) |
| Lesser whitethroat |  |  |  | 1 (0) |  |
| Long-tailed tit |  |  | 1 (1) | 1 (0) |  |
| Reed bunting |  |  | 3 (1) | 2 (0) |  |
| Reed warbler | 1 (0) |  | 2 (2) | 3 (0) |  |
| Robin |  |  | 11 (3) | 7 (2) | 1 (0) |
| Sedge warbler |  |  | 1 (1) | 1 (0) |  |
| Song thrush |  |  |  | 1 (0) |  |
| Whitethroat |  |  | 5 (3) | 7 (2) | 1 (0) |
| Willow warbler |  |  |  | 3 (0) |  |
| Wren | 1 (0) | 1 (0) | 4 (1) | 7 (2) | 1 (0) |
| Yellowhammer |  |  | 1 (0) | 3 (0) |  |

Resistance was highest to vancomycin (n=68, 33%) and tetracycline (n=63, 31%), and lowest to teicoplanin (n=10, 5%) and streptomycin (n=14, 7%; Table S4)

None of Julian day, age, species or site were associated with the presence of *Enterococcus* (Table 7). However, the prevalence of MDR *Enterococcus* declined throughout the season (Table 7). None of diet diversity, migratory status, human habitation or granivorous status were associated with the prevalence of either *Enterococcus* or MDR *Enterococcus* in infected birds, but birds that ate invertebrates had a marginally higher prevalence of MDR *Enterococcus* than birds that did not eat invertebrates (invertebrates: 33.2 ± 3 % [n=190]; no invertebrates: 10.0 ± 10.0 % [n=10]; Table S5).

**Table 7.** Results of binomial generalised linear models testing the effect of environmental variables on the presence of a) *Enterococcus* and b) MDR *Enterococcus* in avian faecal samples.

| a) *Enterococcus* | Dev | df | p |
| --- | --- | --- | --- |
| Age | 1.284 | 1, 239 | 0.257 |
| Species | 17.272 | 15, 238 | 0.303 |
| Julian day | 1.045 | 1, 223 | 0.307 |
| Site | 0.394 | 2, 222 | 0.821 |
| b) MDR *Enterococcus* | Dev | df | P |
| **Julian day** | **7.312** | **1, 199** | **0.007** |
| Age | 1.422 | 1, 198 | 0.233 |
| Species | 23.978 | 21, 197 | 0.294 |
| Site | 1.441 | 2, 176 | 0.487 |

### E. coli

All 259 avian faecal samples were positive for *E. coli*; 153 (59%) of these were resistant to at least three classes of antibiotic (Table S6). Resistance was highest to ampicillin (n=114, 44%) and nalidixic acid (n=106, 41%) and lowest to kanamycin (n=5, 2%) and amikacin (n=6, 2%; Table S6). The presence of MDR *E. coli* did not differ between sites, species, or age classes of bird, or with Julian day (Table 8). Birds with MDR *E. coli* had a higher diet diversity that those without (with MDR *E. coli*: 1.98 ± 0.04 food types [n=150]; without MDR *E. coli*: 1.81 ± 0.05 food types [n=105]); no other host ecological traits were associated with infection by MDR *E. coli* (Table 8).

**Table 8.** Results of binomial generalised linear models testing the effect of a) environmental variables and b) host ecological traits on the presence of MDR *E. coli*.

| a) | Dev | df | p |
| --- | --- | --- | --- |
| Species | 19.360 | 15, 239 | 0.198 |
| Site | 0.671 | 1, 224 | 0.413 |
| Julian day | 1.438 | 1, 222 | 0.231 |
| Age | 0.543 | 1, 221 | 0.461 |
| b) | Dev | df | p |
| **Diet diversity** | **6.354** | **1, 254** | **0.012** |
| Insectivorous | 1.408 | 1, 253 | 0.235 |
| Granivorous | 0.916 | 1, 252 | 0.339 |
| Human habitation | 0.571 | 1, 251 | 0.450 |
| Migratory | 0.037 | 1, 250 | 0.848 |

Terms in bold were retained in the final model

## Discussion

This study is one of very few that investigates multiple pathogens from within the same animal, with many other studies reporting results separately making comparisons difficult. The high levels of antimicrobial resistant bacteria for all species tested (*Campylobacter, Salmonella, Enterococcus* and *E. coli*) is of concern, but perhaps not a surprise given the huge rise in use of AMR seen globally in many different species. Although wildlife AMR is tested much less frequently than companion or livestock AMR, these animals - especially birds due to the long distances which they travel - can provide a good measure of environmental AMR contamination with both AMR bacteria, and resistance genes.

### Salmonella

Comparison of our results with those of previous studies is complicated by the high diversity of bird species, and a large-scale comparison of each individual species would be beneficial to allow for a more meaningful comparison. Many previous studies are carried out in association with farms, potentially to quantify the risk of introduction of the focal pathogen; however, given the routinely high levels of AMR bacteria and *Salmonella* isolated from many farmed livestock species, the results from these previous studies may be highly skewed (De Lucia et al., 2018). In addition, rather than the methods of sampling using mist nets as employed in this study, many focus on birds in rescue shelters, birds such as pigeons that have been culled, or specific focal species such as raptors or large animals such as storks; thus these studies often include only small numbers of passerine birds despite their tendency to be more numerous and widespread. Species with high *Salmonella* prevalence in this study included the Eurasian blackcap (*Sylvia atricapilla),* Eurasian blue tits (*Cyanistes caeruleus*) and house sparrows (*Passer domesticus*). Unfortunately, few previous studies target these species, with a few studies reporting *Salmonella* spp. in the Eurasian Blackcap (Mancini et al. 2020 with a prevalence of 1/8; Foti et al. 2009 (two positives for *Salmonella* spp., but does not say how many blackcaps were tested specifically)), the Eurasian blue tit (Mancini et al. 2020 (one positive from six samples tested); Refsum et al. 2002 (3/14 positive samples) Boonyarittichaikij et al. 2017 (4 of 36 eggs)) and house sparrows (Refsum et al. 2002 (7/31 postiives); Pasquali et al. 2014 (7/11 carcasses); Pennycott, Park, and Mather 2006 who reported 33/50 infected with *Salmonella* spp.).

This is particularly surprising given that studies report that wild bird salmonellosis can pose a risk to human and animal health, and given that a large number of UK homeowners provide supplementary food to garden birds (Robb et al., 2008), this offers a simple yet successful method for introduction of *Salmonella* into the human environment and potentially for infection (Davies et al. 2009; Lawson et al. 2014). Indeed, previous studies have linked wild songbirds to *Salmonella* outbreaks in humans (Patel et al., 2023).

That said, our findings (11.6% *Salmonella* spp. positivity) are in line with, or slightly higher than, other studies, including Krawiec et al. (2015) who reported 6.4% on average for carriage in wild birds in Poland, but this varied dramatically between bird species, with the Eurasian siskin (*Carduelis spinus*) and the greenfinch (*Carduelis chloris*) being around 33% (based on 30 or more samples). A similar prevalence (7.4%) was observed in a study in Croatia (Vlahović et al., 2004) and lower prevalences were observed by Wei et al. (2020) in South Korea (0.93%) and by Konicek et al. (2016) on the Austria-Czech Republic border (2.2%). However, higher prevalences have also been found: for example, in Texas, USA, 17% of wild birds were positive for *Salmonella* spp., (Brobey et al., 2017), as were 13.5% of birds tested in Bangladesh (Card et al., 2023).

The isolation of S. Typhimurium is relatively common among various bird species (Alley et al., 2002; Hughes et al., 2008; Mather et al., 2016; Pennycott et al., 2006), which concurs with our findings. Many of the other strains have also been isolated in previous studies (Antilles et al. 2015; Reche et al. 2003; Kirk, Holmberg, and Jeffrey 2002; Tizard 2004; Refsum et al. 2002; De Lucia et al. 2018; Smith et al. 2023). Interestingly, some of these strains have been isolated from other animals including dogs (Morgan et al., 2023) and livestock (Davies et al. 2004; Snow et al. 2007), suggesting that there may be transmission either from birds to these animals, or vice versa (Horton et al., 2013; Pao et al., 2014). Multi-drug resistance has been reported commonly within *Salmonella* isolates, some of which were from wild birds. Multi drug resistance poses problems for treatment opportunities in infected animals and humans, and thus the clinical shedding of these bacteria is of high importance. Indeed, *Salmonella* spp. isolated from wild birds are commonly MDR, including 86.7% of 15 isolates obtained by Martín-Maldonado et al. (2020).

High levels of tetracycline resistance have been reported previously (Kandir and Öztürk, 2022), with chloramphenicol and ampicillin resistance equally commonly seen in wild bird *Salmonella* isolates (Janecko et al., 2015; Molina-Lopez et al., 2011). The lack of resistance to nalidixic acid seen within this study differs from others which report high levels of resistance (Kandir and Öztürk, 2022; Martín-Maldonado et al., 2020; Troxler et al., 2017), although the reasons for this are unclear.

Risk factor analyses for the carriage of *Salmonella* spp. in wild birds are lacking, as most studies focus on specific populations, such as those admitted to rescue hospitals, or single species. Martin-Maldonado et al. (2020), tested a range of bird species and found that younger birds were more likely to be positive for *Salmonella* than older birds, although we did not find any difference between ages in our study. However, we did find that all intercontinental migrants carrying *Salmonella* were carrying MDR *Salmonella*, which is particularly interesting as these species tend to be reliant on invertebrate food rather than food provided by householders. The review by Blazar, Allard, and Lienau, (2011) suggested that a wide variety of different insects which could act as prey for passerine birds, such as lesser mealworm, *Alphitobius diaperinus* (Panzer) can carry several pathogens, including *Salmonella* spp. or *E. coli* and act as successful vectors, suggesting a potential transmission route. Similarly, dipteran flies, commonly eaten by a range of bird species including migrants, can vector also *Salmonella* spp. (Wales et al., 2010)

### Campylobacter

We found *Campylobacter* spp. presence to differ between species, and many studies have found similar results. However, many focus specifically on *C. jejuni* because this is the most common species to cause disease in humans (Rodrigues et al., 2001). Indeed, Cody et al. (2015) report that wild bird *C. jejuni* are a consistent cause of human disease. Mencía-Gutierrez et al. (2021) report a prevalence of *Campylobacter* spp. in 7.5% in raptors from Spain, with *C. jejuni* making up 88.5% of the isolates and Waldenström et al. (2002) report a prevalence of 21.6% in Sweden, but this was highly variable across different species, and 24.8% prevalence was reported in Italy at a wildlife rescue centre with 94.23% of these being *C. jejuni* and the remained being *C. coli* (Casalino et al., 2022). Similar to the prevalence obtained here, 15.3% was reported in South Korea (Kwon et al., 2017) and ranged from 8.5% to 50% depending on the bird species in Antarctic and sub-Antarctic regions (Johansson et al., 2018). Many of these studies target different bird species, in different areas, and in some cases use different laboratory methodologies, and as such, the results are difficult to compare.

The specific *Campylobacter* species which we isolated are very much in line with other studies, although the most common species seems to be variable. Similar to our study, *C. jejuni* was most common in wild birds associated with a Danish livestock farm (Hald et al., 2015), in an Italian rescue shelter (Casalino et al., 2022), in the mid-Atlantic region of the USA (Keller et al., 2011; Keller and Shriver, 2014), in Northern Poland (Andrzejewska et al., 2022) and from wild birds of prey in Spain (Mencía-Gutiérrez et al., 2021). By contrast, *C. lari* was most commonly found by Waldenstrom et al. (2002) in Sweden, and in the Antarctic peninsula by Johanssen et al. (2018), although we found only a slightly lower prevalence of *C. lari* compared to *C. jejuni*.

Drug resistance levels appear to vary widely across studies, dependent upon area and bird species tested. Variation among the laboratory protocols used also makes direct comparisons difficult. Resistance to tetracycline is common in many studies, and has been reported previously by Casalino et al. (2022) and Du et al. (2019) but tetracycline and amoxicillin resistance was lower in reports by Dudzic et al. (2016) and Marotta et al. (2019). Erythromycin resistance is variable, with low levels reported by Casalino et al. (2022) and Du et al. (2019) whereas Casalino et al. (2022) also reported high levels of resistance to trimethoprim-sulfamethoxazole. By contrast, no drug resistance was found to erythromycin and low resistance was found to amoxicillin by Waldenström et al. (2002) and Kürekci et al. (2021) found no resistance to erythromycin or tetracycline.

We found juvenile birds to be more than twice as likely to be infected by *Campylobacter* spp. than adults, in agreement with Taff et al. (2016), although other studies find no association (e.g. Mencía-Gutiérrez et al. 2021; Jurado-Tarifa et al. 2016). With regards to risk factors from the environment, farmland and animals have been shown to be risk factors for an increased level of *Campylobacter* spp. carriage in wild birds (Hald et al., 2015) and it has been suggested that wild birds may play a role in infection of livestock with *Campylobacter* spp. (Sippy et al., 2012) although other studies suggest that the converse is true (Hughes et al. 2009).

Bird species associated with human habitation had a higher prevalence of *Campylobacter* spp. than those not associated with human habituation, which poses many potential questions, including whether the pathogen comes from human food, or possibly from contact with bird feeders. It also increases the potential risk of transmission from bird to human (or vice versa); indeed, previous epidemics have been linked to wild bird contact (Ahmed and Gulhan, 2022). The risk factors for carriage of *Campylobacter*spp., and the risks which they pose to human health, require further research.

### Enterococcus

Previous studies suggest that the prevalence of *Enterococcus* varies dramatically depending on the bird species tested, geographical area, and the laboratory methodologies used (Kuntz et al., 2004). However, isolates of *Enterococcus* found in wild birds can cause infections in humans (Stępień-Pyśniak et al., 2018).

The prevalence of *Enterococcus* obtained in this study is similar, if slightly higher than that reported in other studies, including 63.3% reported in the Azores archipelago (Santos et al., 2013), 65.8% in Tunisia (Klibi et al., 2015), 66.7% in Poland (Kwit et al., 2023) and 74% in Slovakia (Splichalova et al., 2015). The species isolated in this study are similar to those isolated in other studies, with *E. faecium* being most common in many studies (Yahia et al. 2018; Santos et al. 2013; Radhouani et al. 2012; Klibi et al. 2015; Dolka et al. 2020; Kwit et al. 2023). The other species were isolated in lower numbers in various studies which is also in line with our findings (Yahia et al. 2018; Santos et al. 2013; Klibi et al. 2015; Dolka et al. 2020; Kwit et al. 2023).

Drug resistance is commonly seen in *Enterococcus* spp., with nearly every isolate obtained by Cagnoli et al. (2022) being described as MDR, and this bacterium has become a common indicator of environmental contamination with faecal matter due to its ability to rapidly uptake antimicrobial resistance genes (Suzuki et al., 2012). Resistance to vancomycin is common, especially within *E. faecium* and *E. faecalis* isolates (Reviewed by Wada et al., 2024). Tetracycline resistance was also common in *Enterococcus* isolates in this study, and this is also commonly seen in farm animal isolated *Enterococcus* (Gouliouris et al., 2018) which may offer a potential transmission route for the bacteria, although the direction is unknown. In addition, other studies have reported high levels of resistance of *Enterococcus* to tetracycline (Dec et al., 2020; Kwit et al., 2023; Radhouani et al., 2012; Santos et al., 2013). Surprisingly in this study, resistance to teicoplanin and streptomycin were low, which is in contrast to the results reported by Dec et al. (2020), although other studies support the low resistance finding to these antibiotics including Radhouani et al. (2012), Kwit et al. (2023) and Santos et al. (2013). Previous studies have suggested that the level of antibiotic resistance genes is not consistent across the year, with crows shown to carry lower levels of antibiotic resistance in the summer compared to ducks and gulls, and the authors attribute this to seasonal variation in food resources due to winter foraging in waste disposal areas and highly populated areas compared to summer where seeds and grain make up more of the diets (Dolejska and Literak, 2019; Zhao et al., 2020). This concurs with our findings of a decline in the prevalence of MDR *Enterococcus* through the season: it is likely that temperature, humidity and density of animals will have an impact on the carriage and transmission of antimicrobial resistance genes (Bharathi et al., 2019).

Birds with a wider dietary range may tend to carry more pathogens (Oravcova et al., 2014; Radhouani et al., 2012), which concurs with our finding of a tendency for insectivorous birds to carry a higher prevalence of *Enterococcus* than other birds. Insects have been shown to be a common carrier of *Enterococcus* spp. as well as other bacteria, and this may allow for a route of transmission to birds (Reviewed by Rawat et al., 2023). In addition, *Enterococcus* and other antimicrobial resistant bacteria have been isolated from caterpillars, which act as one of the major food sources for many insectivorous birds and may allow for a transmission route (Huff et al., 2020).

### E. coli

*E. coli* is very commonly isolated from the faecal samples of many animals and is often used to assess antimicrobial resistance. Consequently, many studies have been carried out in birds, but results depend on the geographic areas and the bird species tested. This is epitomised by the study by Stedt et al. (2014) who reported a variation in *E. coli* resistance in gulls in Europe varying from 61.2% in Spain to 8.3% in Denmark. In the Azores archipelago, AMR within *E. coli* isolates was shown to be 24.3% (Santos et al., 2013), but increases to values of 63% in Turkey (Ahmed and Gulhan, 2024). Resistance levels vary, although similarly to our study, resistance to ampicillin tends to be common (Guenther et al., 2010; Merkeviciene et al., 2018; Ong et al., 2020; Santos et al., 2013; Skarżyńska et al., 2021), although other studies such as that conducted in Poland by Nowaczek et al. (2021) report a lower prevalence at 28.1% and 16.7% reported by Yuan et al. (2021). Total resistance to ampicillin has also been reported (Rybak et al., 2022). In addition, nalidixic acid resistance also seems common in wild birds in various parts of the world (Ong et al., 2020; Ramey et al., 2018; Skarżyńska et al., 2021). However, the resistance to amikacin seems variable with Santos et al. (2013) and Yuan et al. (2021) reporting a low level of resistance to this antibiotic, or in some cases, no resistance at all (Ong et al. 2020; Merkeviciene et al. 2018; Smith et al. 2019) whereas Prandi et al. (2023) report amikacin resistance of 17.9%. Kanamycin resistance again varies among wild bird *E. coli* isolates, with 18.7% resistance reported by Nowaczek et al. (2021) but much higher resistance of 38% observed in birds in Turkey (Ahmed and Gulhan, 2022).

Levels of MDR also tend to vary, with Prandi et al. (2023) reporting 39.6% of *E. coli* isolates being resistant to three or more antimicrobials in Italy, 31.2 % MDR observed in Poland (Nowaczek et al., 2021), 38% in Brazil (Batalha De Jesus et al., 2019), 33.5% in Lithuania (Merkeviciene et al., 2018) and 38.6% in Poland (Skarżyńska et al., 2021). Similar to this study, Yuan et al. (2021) found 61.9% of 118 isolates which were classed as MDR. Total MDR (i.e. 100% of isolated bacteria) was observed in some villages in Malaysia, but the levels varied by area (Mohamed et al., 2022). Interestingly, our findings suggested a higher diet diversity in individuals carrying MDR *E. coli* compared to those carrying non-MDR *E. coli*, suggesting that exposure to *E. coli* in multiple food types may increase the likelihood of MDR (Radhouani et al., 2012).

## Conclusion

The high level of pathogen carriage observed within this study from birds within the UK acts as a timely reminder of the risks which bird contact and bird faecal matter may pose, and the impacts that land management can have on wildlife. Although contact with wild birds is generally limited, risks may be posed from bird feeders, or through indiscriminate defecation in urban or suburban areas leading to environmental contamination. This in turn may lead to infections of other animals such as companion animals or livestock and could potentially enter the food chain leading to zoonotic risks. Although not tested in this study, the presence of antimicrobial resistance genes are also likely to pose potential risks to humans and animals. It is crucial that further research tests potential mechanisms of reducing levels of MDR bacteria in wildlife, potentially through increased hygiene of supplementary food resources.

## Ethics

All birds from which samples were collected were caught as part of standard bird ringing activities under a British Trust for Ornithology ringing licence to JCD. This study received ethical approval from the University of Lincoln Animal Ethics Committee, reference LEAS3818.

## Data Availability

Data will be made available on FigShare on acceptance. For review purposes, code is available here:

https://figshare.com/s/eafb48a1091c3be68d4d and datasets are available here:

https://figshare.com/s/816e886090183b7b68ac (full dataset),

https://figshare.com/s/bb3666087b5c67451c89 (full dataset excluding species with n<4),

https://figshare.com/s/3e5342211d9188cd2faf (all *Campylobacter* positive samples),

https://figshare.com/s/bc3987ac2b3a0a958286 (*Campylobacter* positive samples, excluding host species with n<4),

https://figshare.com/s/aa188386eaaf96667ddd (all *Salmonella* positive samples), and

https://figshare.com/s/056f3ff48d5878e7ec31 (all *Enterococcus* positive samples).

The following DOIs have been reserved and will be activated upon manuscript acceptance.

Data are available through FigShare at the following DOIs. Analysis code is available at 10.6084/m9.figshare.26160301, with the full dataset available at 10.6084/m9.figshare.26160361, the full dataset excluding species with n<4 available at 10.6084/m9.figshare.26160343, all *Campylobacter* positive samples available at 10.6084/m9.figshare.26160334, *Campylobacter* positive samples excluding host species with n<4 available at 10.6084/m9.figshare.26160349, all Salmonella positive samples available at 10.6084/m9.figshare.26160352, and all *Enterococcus* positive samples available at 10.6084/m9.figshare.26160358.

## Author Contributions

JCD and SRC conceived the study. JCD collected the samples, SRC conducted laboratory analysis. JCD conducted statistical analysis, and SRC and JCD wrote the original draft and reviewed and edited the manuscript.

## Supplementary data

**Table S1.** Antibiotic susceptibility of *Salmonella* isolates recovered from passerine faecal samples, split by host species.

| Species (no. positive) | AK | AMC | AMP | CRO | C | CN | NA | NX | SXT | TE |
| --- | --- | --- | --- | --- | --- | --- | --- | --- | --- | --- |
| Blackbird (n=2) |  |  |  |  |  |  |  |  |  |  |
| Resistant | 0 | 0 | 0 | 0 | 2 | 0 | 0 | 0 | 0 | 1 |
| Intermediate | 0 | 0 | 0 | 1 | 0 | 0 | 0 | 1 | 1 | 1 |
| Susceptible | 2 | 2 | 2 | 1 | 0 | 2 | 2 | 1 | 1 | 0 |
| Blackcap (n=5) |  |  |  |  |  |  |  |  |  |  |
| Resistant | 0 | 2 | 3 | 4 | 3 | 0 | 0 | 0 | 3 | 5 |
| Intermediate | 0 | 1 | 0 | 0 | 0 | 2 | 1 | 0 | 1 | 0 |
| Susceptible | 5 | 2 | 2 | 1 | 2 | 3 | 4 | 5 | 1 | 0 |
| Blue tit (n=3) |  |  |  |  |  |  |  |  |  |  |
| Resistant | 0 | 0 | 2 | 1 | 1 | 0 | 0 | 0 | 1 | 1 |
| Intermediate | 0 | 1 | 0 | 1 | 0 | 0 | 0 | 0 | 0 | 1 |
| Susceptible | 3 | 2 | 1 | 1 | 2 | 3 | 3 | 3 | 2 | 1 |
| Chaffinch (n=2) |  |  |  |  |  |  |  |  |  |  |
| Resistant | 0 | 1 | 0 | 0 | 0 | 0 | 0 | 0 | 1 | 2 |
| Intermediate | 0 | 0 | 0 | 0 | 1 | 0 | 0 | 0 | 1 | 0 |
| Susceptible | 2 | 1 | 2 | 2 | 1 | 2 | 2 | 2 | 0 | 0 |
| Chiffchaff (n=1) |  |  |  |  |  |  |  |  |  |  |
| Resistant | 0 | 1 | 0 | 1 | 1 | 0 | 0 | 0 | 0 | 1 |
| Intermediate | 0 | 0 | 0 | 0 | 0 | 0 | 0 | 0 | 1 | 0 |
| Susceptible | 1 | 0 | 1 | 0 | 0 | 1 | 1 | 1 | 0 | 0 |
| Dunnock (n=2) |  |  |  |  |  |  |  |  |  |  |
| Resistant | 0 | 1 | 1 | 1 | 0 | 0 | 0 | 0 | 1 | 1 |
| Intermediate | 0 | 0 | 0 | 1 | 0 | 0 | 1 | 0 | 0 | 0 |
| Susceptible | 2 | 1 | 1 | 0 | 2 | 2 | 1 | 2 | 1 | 1 |
| Goldfinch (n=2) |  |  |  |  |  |  |  |  |  |  |
| Resistant | 0 | 0 | 2 | 1 | 1 | 0 | 0 | 0 | 1 | 1 |
| Intermediate | 0 | 0 | 0 | 1 | 0 | 0 | 0 | 0 | 0 | 1 |
| Susceptible | 2 | 2 | 0 | 0 | 1 | 2 | 2 | 2 | 1 | 0 |
| House sparrow (n=8) |  |  |  |  |  |  |  |  |  |  |
| Resistant | 1 | 3 | 4 | 4 | 6 | 0 | 1 | 0 | 4 | 5 |
| Intermediate | 0 | 0 | 0 | 2 | 1 | 2 | 0 | 2 | 2 | 0 |
| Susceptible | 7 | 5 | 4 | 2 | 1 | 6 | 7 | 6 | 2 | 3 |
| Robin (n=2) |  |  |  |  |  |  |  |  |  |  |
| Resistant | 0 | 0 | 0 | 0 | 2 | 0 | 0 | 0 | 0 | 2 |
| Intermediate | 0 | 0 | 0 | 2 | 0 | 0 | 0 | 0 | 0 | 0 |
| Susceptible | 2 | 2 | 2 | 0 | 0 | 2 | 2 | 2 | 2 | 0 |
| Whitethroat (n=1) |  |  |  |  |  |  |  |  |  |  |
| Resistant | 0 | 1 | 1 | 1 | 1 | 0 | 0 | 0 | 0 | 0 |
| Intermediate | 0 | 0 | 0 | 0 | 0 | 0 | 0 | 0 | 0 | 0 |
| Susceptible | 1 | 0 | 0 | 0 | 0 | 1 | 1 | 1 | 1 | 1 |
| Wren (n=2) |  |  |  |  |  |  |  |  |  |  |
| Resistant | 0 | 1 | 2 | 1 | 0 | 1 | 0 | 0 | 2 | 2 |
| Intermediate | 0 | 0 | 0 | 0 | 0 | 0 | 1 | 0 | 0 | 0 |
| Susceptible | 2 | 1 | 0 | 1 | 2 | 1 | 1 | 2 | 0 | 0 |
| Total (n=30) |  |  |  |  |  |  |  |  |  |  |
| Resistant | 1 | 10 | 15 | 14 | 17 | 1 | 1 | 0 | 13 | 21 |
| Intermediate | 0 | 2 | 0 | 8 | 2 | 4 | 3 | 3 | 6 | 3 |
| Susceptible | 29 | 18 | 15 | 8 | 11 | 25 | 26 | 27 | 11 | 6 |

AK: Amikacin (30 mg/ml); AMC: Amoxicillin-Clavulanic acid (20 mg/ml); AMP: Ampicillin (10 mg/ml); CRO: Ceftriaxone (30 mg); C: Chloramphenicol (30 mg/ml); CN: Gentamycin (10 mg/ml); NA: Nalidixic acid (30 mg/ml); NX: Norfloxacin (10 mg/ml); SXT: Sulfamethoxazole-trimethoprim (25 mg/ml); TE: Tetracycline (30 mg/ml)

**Table S2.** Results of binomial generalised linear models testing the effect of environmental variables on the presence of a) *Salmonella* and b) MDR *Salmonella* in avian faecal samples. Species was excluded as a term for b) due to small sample sizes.

| a) *Salmonella* | Dev | df | p |
| --- | --- | --- | --- |
| Species | 21.700 | 15, 239 | 0.116 |
| Julian day | 1.500 | 1, 223 | 0.221 |
| Age | 0.006 | 1, 222 | 0.938 |
| Site | 0.143 | 2, 220 | 0.931 |
| b) MDR *Salmonella* | Dev | df | p |
| Age | 0.731 | 1, 28 | 0.393 |
| Julian day | 0.161 | 1, 27 | 0.688 |
| Site | 0.145 | 2, 26 | 0.930 |

No terms were retained in the final models.

**Table S3.** Antibiotic susceptibility of *Campylobacter* isolates recovered from passerine faecal samples.

| Species (no. positive) | AZM | AMX | C | CIP | CN | ENR | E | NA | SXT | TE |
| --- | --- | --- | --- | --- | --- | --- | --- | --- | --- | --- |
| Blackbird (n=3) |  |  |  |  |  |  |  |  |  |  |
| Resistant | 3 | 1 | 2 | 2 | 2 | 0 | 1 | 0 | 0 | 1 |
| Intermediate | 0 | 1 | 0 | 1 | 0 | 0 | 0 | 2 | 0 | 1 |
| Susceptible | 0 | 1 | 1 | 0 | 1 | 3 | 2 | 1 | 3 | 1 |
| Blackcap (n=3) |  |  |  |  |  |  |  |  |  |  |
| Resistant | 2 | 1 | 0 | 2 | 1 | 0 | 2 | 1 | 0 | 2 |
| Intermediate | 0 | 1 | 1 | 1 | 0 | 0 | 0 | 1 | 1 | 1 |
| Susceptible | 1 | 1 | 2 | 0 | 2 | 3 | 1 | 1 | 2 | 0 |
| Blue tit (n=5) |  |  |  |  |  |  |  |  |  |  |
| Resistant | 2 | 2 | 0 | 3 | 4 | 2 | 2 | 5 | 1 | 3 |
| Intermediate | 1 | 2 | 2 | 1 | 0 | 2 | 1 | 0 | 0 | 0 |
| Susceptible | 2 | 1 | 3 | 1 | 1 | 1 | 2 | 0 | 4 | 2 |
| Chaffinch (n=3) |  |  |  |  |  |  |  |  |  |  |
| Resistant | 3 | 3 | 0 | 2 | 1 | 0 | 3 | 0 | 0 | 3 |
| Intermediate | 0 | 0 | 1 | 0 | 1 | 0 | 0 | 1 | 1 | 0 |
| Susceptible | 0 | 0 | 2 | 1 | 1 | 3 | 0 | 2 | 2 | 0 |
| Chiffchaff (n=1) |  |  |  |  |  |  |  |  |  |  |
| Resistant | 1 | 0 | 1 | 0 | 0 | 0 | 0 | 1 | 0 | 1 |
| Intermediate | 0 | 0 | 0 | 0 | 0 | 0 | 0 | 0 | 0 | 0 |
| Susceptible | 0 | 1 | 0 | 1 | 1 | 1 | 1 | 0 | 1 | 0 |
| Goldfinch (n=1) |  |  |  |  |  |  |  |  |  |  |
| Resistant | 1 | 1 | 1 | 0 | 0 | 0 | 1 | 0 | 0 | 1 |
| Intermediate | 0 | 0 | 0 | 0 | 0 | 0 | 0 | 0 | 0 | 0 |
| Susceptible | 0 | 0 | 0 | 1 | 1 | 1 | 0 | 1 | 1 | 0 |
| Great tit (n=1) |  |  |  |  |  |  |  |  |  |  |
| Resistant | 1 | 1 | 0 | 0 | 0 | 1 | 1 | 0 | 0 | 0 |
| Intermediate | 0 | 0 | 0 | 0 | 1 | 0 | 0 | 0 | 0 | 0 |
| Susceptible | 0 | 0 | 1 | 1 | 0 | 0 | 0 | 1 | 1 | 1 |
| Greenfinch (n=1) |  |  |  |  |  |  |  |  |  |  |
| Resistant | 1 | 1 | 0 | 1 | 0 | 0 | 1 | 0 | 0 | 1 |
| Intermediate | 0 | 0 | 0 | 0 | 0 | 0 | 0 | 0 | 0 | 0 |
| Susceptible | 0 | 0 | 1 | 0 | 1 | 1 | 0 | 1 | 1 | 0 |
| House sparrow (n=19) |  |  |  |  |  |  |  |  |  |  |
| Resistant | 5 | 12 | 4 | 5 | 8 | 3 | 10 | 1 | 4 | 12 |
| Intermediate | 6 | 4 | 1 | 2 | 3 | 4 | 3 | 8 | 5 | 6 |
| Susceptible | 8 | 3 | 14 | 12 | 8 | 12 | 6 | 10 | 10 | 1 |
| Reed warbler (n=1) |  |  |  |  |  |  |  |  |  |  |
| Resistant | 1 | 1 | 0 | 0 | 1 | 1 | 1 | 0 | 0 | 0 |
| Intermediate | 0 | 0 | 0 | 0 | 0 | 0 | 0 | 1 | 0 | 0 |
| Susceptible | 0 | 0 | 1 | 1 | 0 | 0 | 0 | 0 | 1 | 1 |
| Robin (n=4) |  |  |  |  |  |  |  |  |  |  |
| Resistant | 0 | 1 | 1 | 2 | 2 | 2 | 1 | 3 | 1 | 2 |
| Intermediate | 1 | 2 | 1 | 2 | 1 | 1 | 1 | 0 | 0 | 0 |
| Susceptible | 3 | 1 | 2 | 0 | 1 | 1 | 2 | 1 | 3 | 2 |
| Song thrush (n=1) |  |  |  |  |  |  |  |  |  |  |
| Resistant | 0 | 1 | 0 | 0 | 0 | 0 | 1 | 0 | 0 | 1 |
| Intermediate | 0 | 0 | 0 | 0 | 1 | 0 | 0 | 0 | 0 | 0 |
| Susceptible | 1 | 0 | 1 | 1 | 0 | 1 | 0 | 1 | 1 | 0 |
| Whitethroat (n=3) |  |  |  |  |  |  |  |  |  |  |
| Resistant | 0 | 2 | 2 | 2 | 1 | 0 | 2 | 1 | 1 | 1 |
| Intermediate | 0 | 1 | 1 | 1 | 1 | 1 | 0 | 1 | 0 | 2 |
| Susceptible | 3 | 0 | 0 | 0 | 1 | 2 | 1 | 1 | 2 | 0 |
| Willow warbler (n=1) |  |  |  |  |  |  |  |  |  |  |
| Resistant | 0 | 1 | 0 | 1 | 0 | 0 | 1 | 0 | 0 | 0 |
| Intermediate | 0 | 0 | 0 | 0 | 1 | 0 | 0 | 1 | 0 | 0 |
| Susceptible | 1 | 0 | 1 | 0 | 0 | 1 | 0 | 0 | 1 | 1 |
| Wren (n=2) |  |  |  |  |  |  |  |  |  |  |
| Resistant | 1 | 2 | 0 | 2 | 1 | 0 | 2 | 0 | 0 | 1 |
| Intermediate | 1 | 0 | 0 | 0 | 0 | 0 | 0 | 0 | 1 | 1 |
| Susceptible | 0 | 0 | 2 | 0 | 1 | 2 | 0 | 2 | 1 | 0 |
| Total (n=49) |  |  |  |  |  |  |  |  |  |  |
| Resistant | 21 | 30 | 11 | 22 | 21 | 9 | 29 | 12 | 7 | 29 |
| Intermediate | 9 | 11 | 7 | 8 | 9 | 8 | 5 | 15 | 8 | 11 |
| Susceptible | 19 | 8 | 31 | 19 | 19 | 32 | 15 | 22 | 34 | 9 |

AZM: Azithromycin (15 mg); AMX: Amoxicillin (30 mg); C: Chloramphenicol (30 mg); CIP: Ciprofloxacin (5 mg); CN: Gentamicin (10 mg); ENR: Enrofloxacin (5 mg); E: Erythromycin (15 mg); NA: Nalidixic acid (30 mg); SXT: Trimethoprim-sulfamethoxazole (25 mg); TE: Tetracycline 173 (30 mg)

**Table S4.**
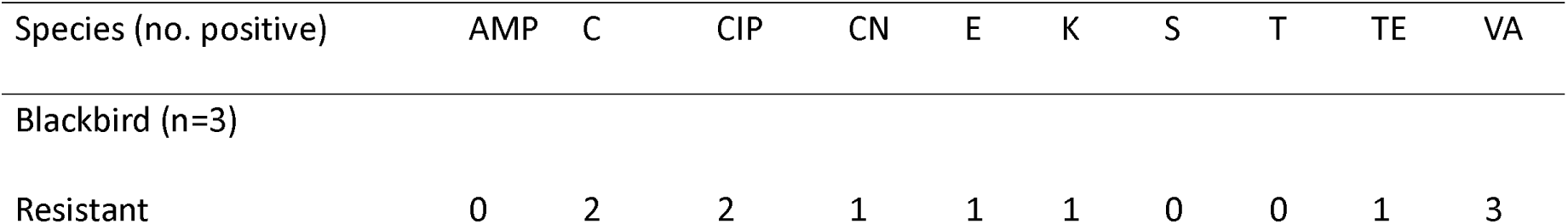
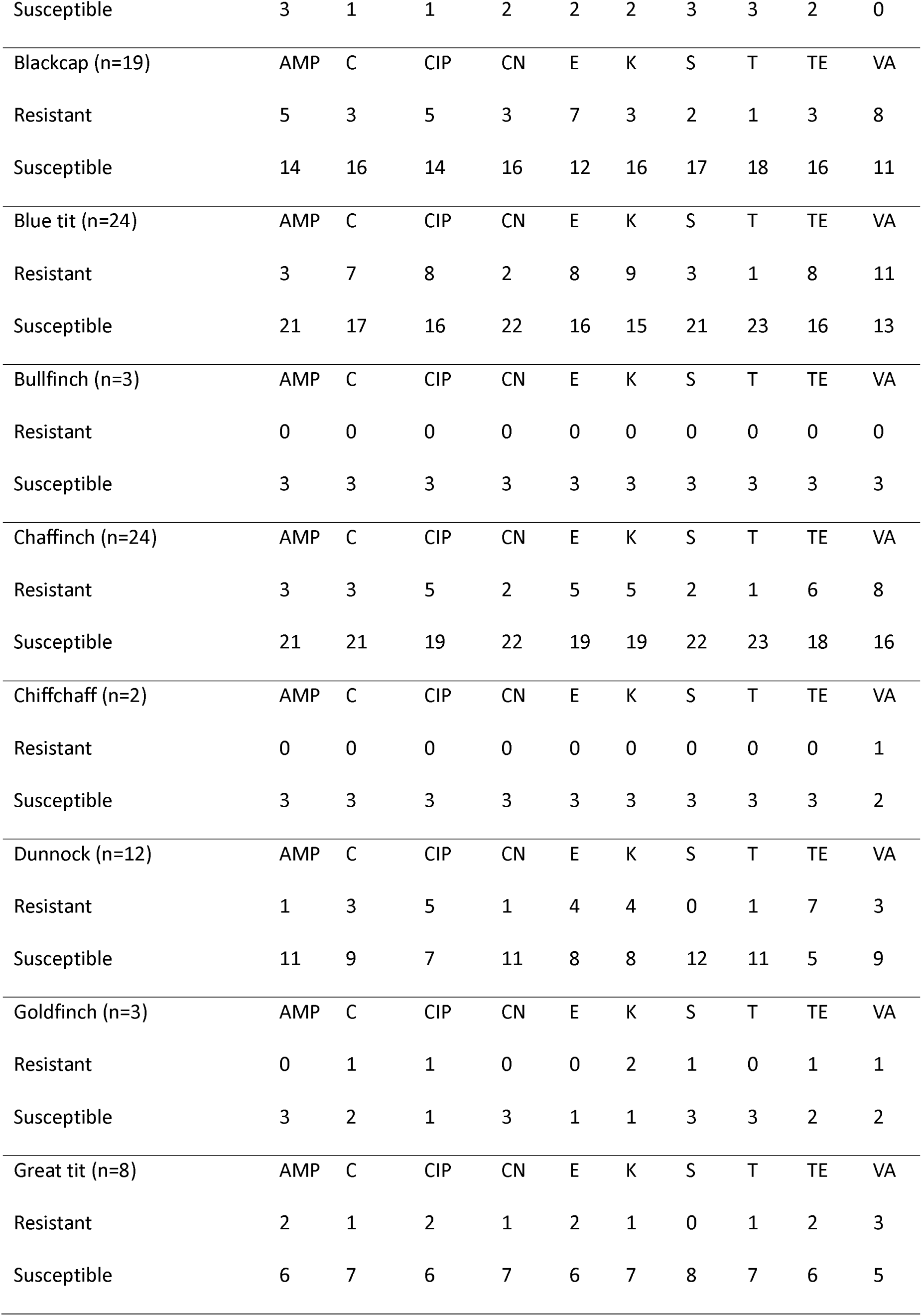
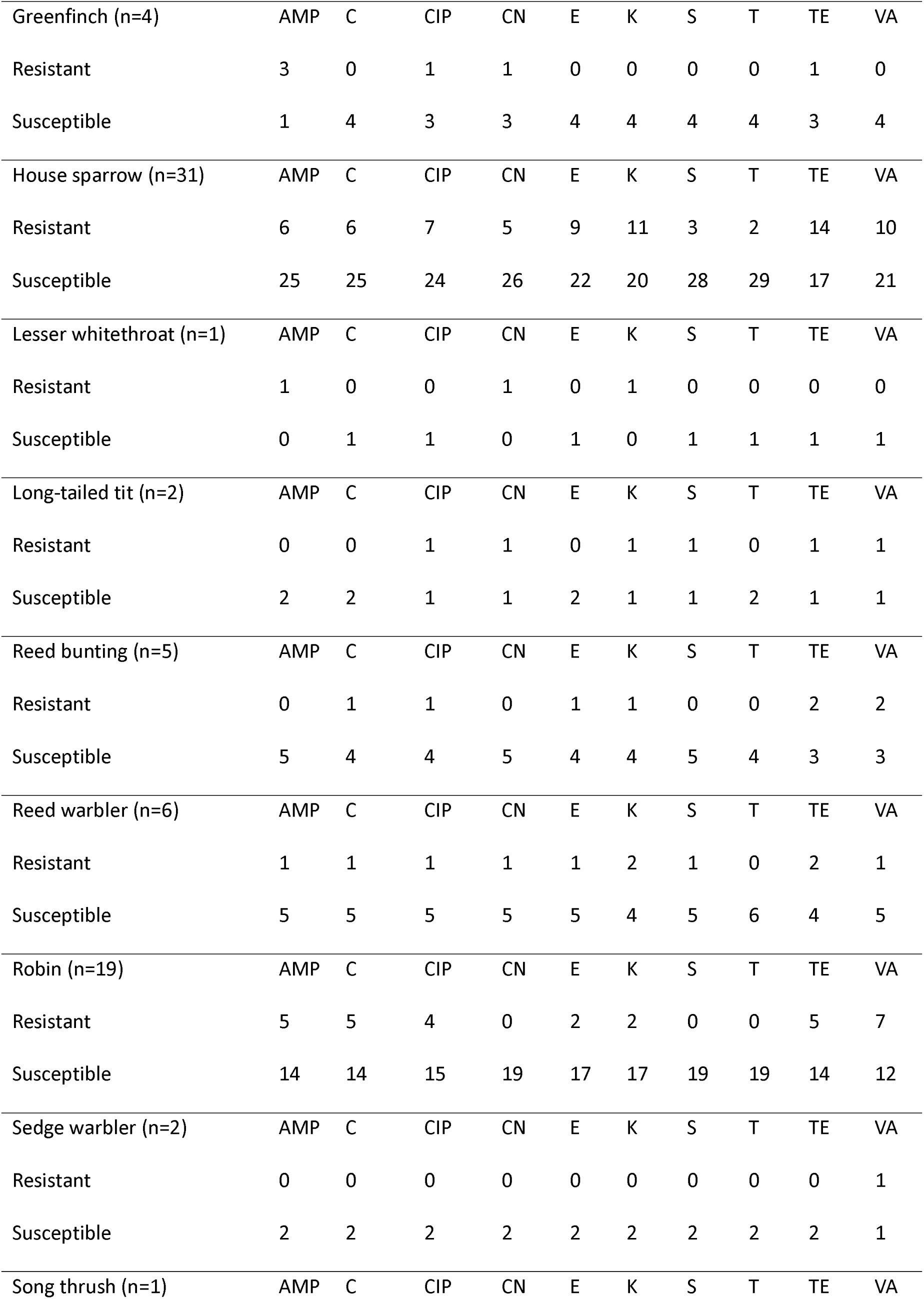
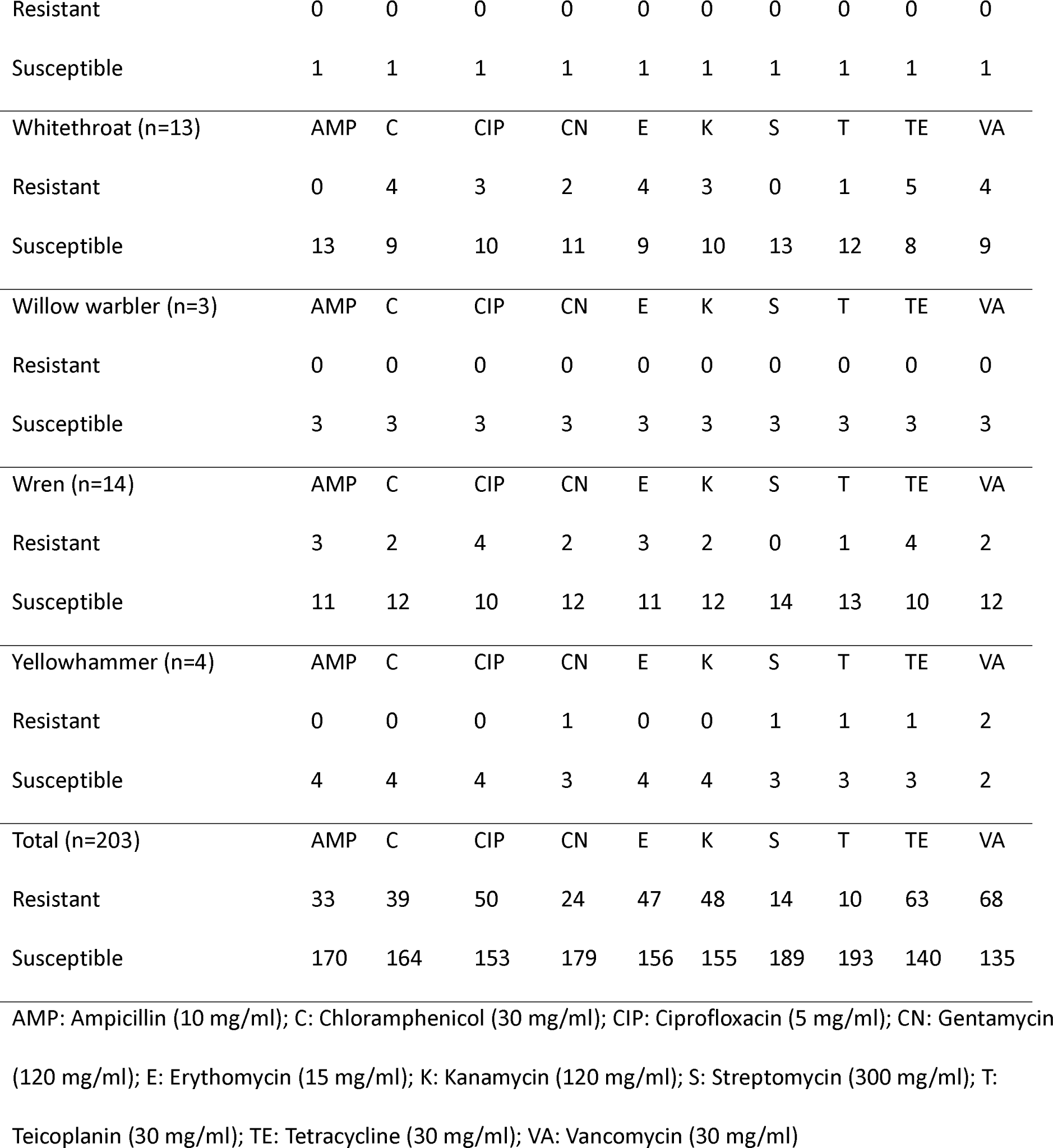
Antibiotic susceptibility of *Enterococcus* isolates recovered from passerine faecal samples.

| Species (no. positive) | AMP | C | CIP | CN | E | K | S | T | TE | VA |
| --- | --- | --- | --- | --- | --- | --- | --- | --- | --- | --- |
| Blackbird (n=3) |  |  |  |  |  |  |  |  |  |  |
| Resistant | 0 | 2 | 2 | 1 | 1 | 1 | 0 | 0 | 1 | 3 |
| Susceptible | 3 | 1 | 1 | 2 | 2 | 2 | 3 | 3 | 2 | 0 |
| Blackcap (n=19) | AMP | C | CIP | CN | E | K | S | T | TE | VA |
| Resistant | 5 | 3 | 5 | 3 | 7 | 3 | 2 | 1 | 3 | 8 |
| Susceptible | 14 | 16 | 14 | 16 | 12 | 16 | 17 | 18 | 16 | 11 |
| Blue tit (n=24) | AMP | C | CIP | CN | E | K | S | T | TE | VA |
| Resistant | 3 | 7 | 8 | 2 | 8 | 9 | 3 | 1 | 8 | 11 |
| Susceptible | 21 | 17 | 16 | 22 | 16 | 15 | 21 | 23 | 16 | 13 |
| Bullfinch (n=3) | AMP | C | CIP | CN | E | K | S | T | TE | VA |
| Resistant | 0 | 0 | 0 | 0 | 0 | 0 | 0 | 0 | 0 | 0 |
| Susceptible | 3 | 3 | 3 | 3 | 3 | 3 | 3 | 3 | 3 | 3 |
| Chaffinch (n=24) | AMP | C | CIP | CN | E | K | S | T | TE | VA |
| Resistant | 3 | 3 | 5 | 2 | 5 | 5 | 2 | 1 | 6 | 8 |
| Susceptible | 21 | 21 | 19 | 22 | 19 | 19 | 22 | 23 | 18 | 16 |
| Chiffchaff (n=2) | AMP | C | CIP | CN | E | K | S | T | TE | VA |
| Resistant | 0 | 0 | 0 | 0 | 0 | 0 | 0 | 0 | 0 | 1 |
| Susceptible | 3 | 3 | 3 | 3 | 3 | 3 | 3 | 3 | 3 | 2 |
| Dunnock (n=12) | AMP | C | CIP | CN | E | K | S | T | TE | VA |
| Resistant | 1 | 3 | 5 | 1 | 4 | 4 | 0 | 1 | 7 | 3 |
| Susceptible | 11 | 9 | 7 | 11 | 8 | 8 | 12 | 11 | 5 | 9 |
| Goldfinch (n=3) | AMP | C | CIP | CN | E | K | S | T | TE | VA |
| Resistant | 0 | 1 | 1 | 0 | 0 | 2 | 1 | 0 | 1 | 1 |
| Susceptible | 3 | 2 | 1 | 3 | 1 | 1 | 3 | 3 | 2 | 2 |
| Great tit (n=8) | AMP | C | CIP | CN | E | K | S | T | TE | VA |
| Resistant | 2 | 1 | 2 | 1 | 2 | 1 | 0 | 1 | 2 | 3 |
| Susceptible | 6 | 7 | 6 | 7 | 6 | 7 | 8 | 7 | 6 | 5 |
| Greenfinch (n=4) | AMP | C | CIP | CN | E | K | S | T | TE | VA |
| Resistant | 3 | 0 | 1 | 1 | 0 | 0 | 0 | 0 | 1 | 0 |
| Susceptible | 1 | 4 | 3 | 3 | 4 | 4 | 4 | 4 | 3 | 4 |
| House sparrow (n=31) | AMP | C | CIP | CN | E | K | S | T | TE | VA |
| Resistant | 6 | 6 | 7 | 5 | 9 | 11 | 3 | 2 | 14 | 10 |
| Susceptible | 25 | 25 | 24 | 26 | 22 | 20 | 28 | 29 | 17 | 21 |
| Lesser whitethroat (n=1) | AMP | C | CIP | CN | E | K | S | T | TE | VA |
| Resistant | 1 | 0 | 0 | 1 | 0 | 1 | 0 | 0 | 0 | 0 |
| Susceptible | 0 | 1 | 1 | 0 | 1 | 0 | 1 | 1 | 1 | 1 |
| Long-tailed tit (n=2) | AMP | C | CIP | CN | E | K | S | T | TE | VA |
| Resistant | 0 | 0 | 1 | 1 | 0 | 1 | 1 | 0 | 1 | 1 |
| Susceptible | 2 | 2 | 1 | 1 | 2 | 1 | 1 | 2 | 1 | 1 |
| Reed bunting (n=5) | AMP | C | CIP | CN | E | K | S | T | TE | VA |
| Resistant | 0 | 1 | 1 | 0 | 1 | 1 | 0 | 0 | 2 | 2 |
| Susceptible | 5 | 4 | 4 | 5 | 4 | 4 | 5 | 4 | 3 | 3 |
| Reed warbler (n=6) | AMP | C | CIP | CN | E | K | S | T | TE | VA |
| Resistant | 1 | 1 | 1 | 1 | 1 | 2 | 1 | 0 | 2 | 1 |
| Susceptible | 5 | 5 | 5 | 5 | 5 | 4 | 5 | 6 | 4 | 5 |
| Robin (n=19) | AMP | C | CIP | CN | E | K | S | T | TE | VA |
| Resistant | 5 | 5 | 4 | 0 | 2 | 2 | 0 | 0 | 5 | 7 |
| Susceptible | 14 | 14 | 15 | 19 | 17 | 17 | 19 | 19 | 14 | 12 |
| Sedge warbler (n=2) | AMP | C | CIP | CN | E | K | S | T | TE | VA |
| Resistant | 0 | 0 | 0 | 0 | 0 | 0 | 0 | 0 | 0 | 1 |
| Susceptible | 2 | 2 | 2 | 2 | 2 | 2 | 2 | 2 | 2 | 1 |
| Song thrush (n=1) | AMP | C | CIP | CN | E | K | S | T | TE | VA |
| Resistant | 0 | 0 | 0 | 0 | 0 | 0 | 0 | 0 | 0 | 0 |
| Susceptible | 1 | 1 | 1 | 1 | 1 | 1 | 1 | 1 | 1 | 1 |
| Whitethroat (n=13) | AMP | C | CIP | CN | E | K | S | T | TE | VA |
| Resistant | 0 | 4 | 3 | 2 | 4 | 3 | 0 | 1 | 5 | 4 |
| Susceptible | 13 | 9 | 10 | 11 | 9 | 10 | 13 | 12 | 8 | 9 |
| Willow warbler (n=3) | AMP | C | CIP | CN | E | K | S | T | TE | VA |
| Resistant | 0 | 0 | 0 | 0 | 0 | 0 | 0 | 0 | 0 | 0 |
| Susceptible | 3 | 3 | 3 | 3 | 3 | 3 | 3 | 3 | 3 | 3 |
| Wren (n=14) | AMP | C | CIP | CN | E | K | S | T | TE | VA |
| Resistant | 3 | 2 | 4 | 2 | 3 | 2 | 0 | 1 | 4 | 2 |
| Susceptible | 11 | 12 | 10 | 12 | 11 | 12 | 14 | 13 | 10 | 12 |
| Yellowhammer (n=4) | AMP | C | CIP | CN | E | K | S | T | TE | VA |
| Resistant | 0 | 0 | 0 | 1 | 0 | 0 | 1 | 1 | 1 | 2 |
| Susceptible | 4 | 4 | 4 | 3 | 4 | 4 | 3 | 3 | 3 | 2 |
| Total (n=203) | AMP | C | CIP | CN | E | K | S | T | TE | VA |
| Resistant | 33 | 39 | 50 | 24 | 47 | 48 | 14 | 10 | 63 | 68 |
| Susceptible | 170 | 164 | 153 | 179 | 156 | 155 | 189 | 193 | 140 | 135 |

AMP: Ampicillin (10 mg/ml); C: Chloramphenicol (30 mg/ml); CIP: Ciprofloxacin (5 mg/ml); CN: Gentamycin (120 mg/ml); E: Erythromycin (15 mg/ml); K: Kanamycin (120 mg/ml); S: Streptomycin (300 mg/ml); T: Teicoplanin (30 mg/ml); TE: Tetracycline (30 mg/ml); VA: Vancomycin (30 mg/ml)

**Table S5.** Results of binomial generalised linear models testing the effect of host ecological traits on the presence of a) *Enterococcus* and b) MDR *Enterococcus* in avian faecal samples.

| a) *Enterococcus* | Dev | df | p |
| --- | --- | --- | --- |
| Granivorous | 1.624 | 1, 254 | 0.203 |
| Human habitation | 1.976 | 1, 253 | 0.160 |
| Insectivorous | 0.939 | 1, 252 | 0.332 |
| Migratory | 0.589 | 1, 251 | 0.443 |
| Diet diversity | 0.025 | 1, 250 | 0.873 |

| b) MDR *Enterococcus* | Dev | df | p |
| --- | --- | --- | --- |
| *Insectivorous* | 2.836 | 1, 199 | 0.092 |
| Human habitation | 1.861 | 1, 198 | 0.173 |
| Diet diversity | 0.388 | 1, 197 | 0.533 |
| Granivorous | 0.213 | 1, 196 | 0.645 |
| Migratory | 0.021 | 1, 195 | 0.884 |

**Table S6.**
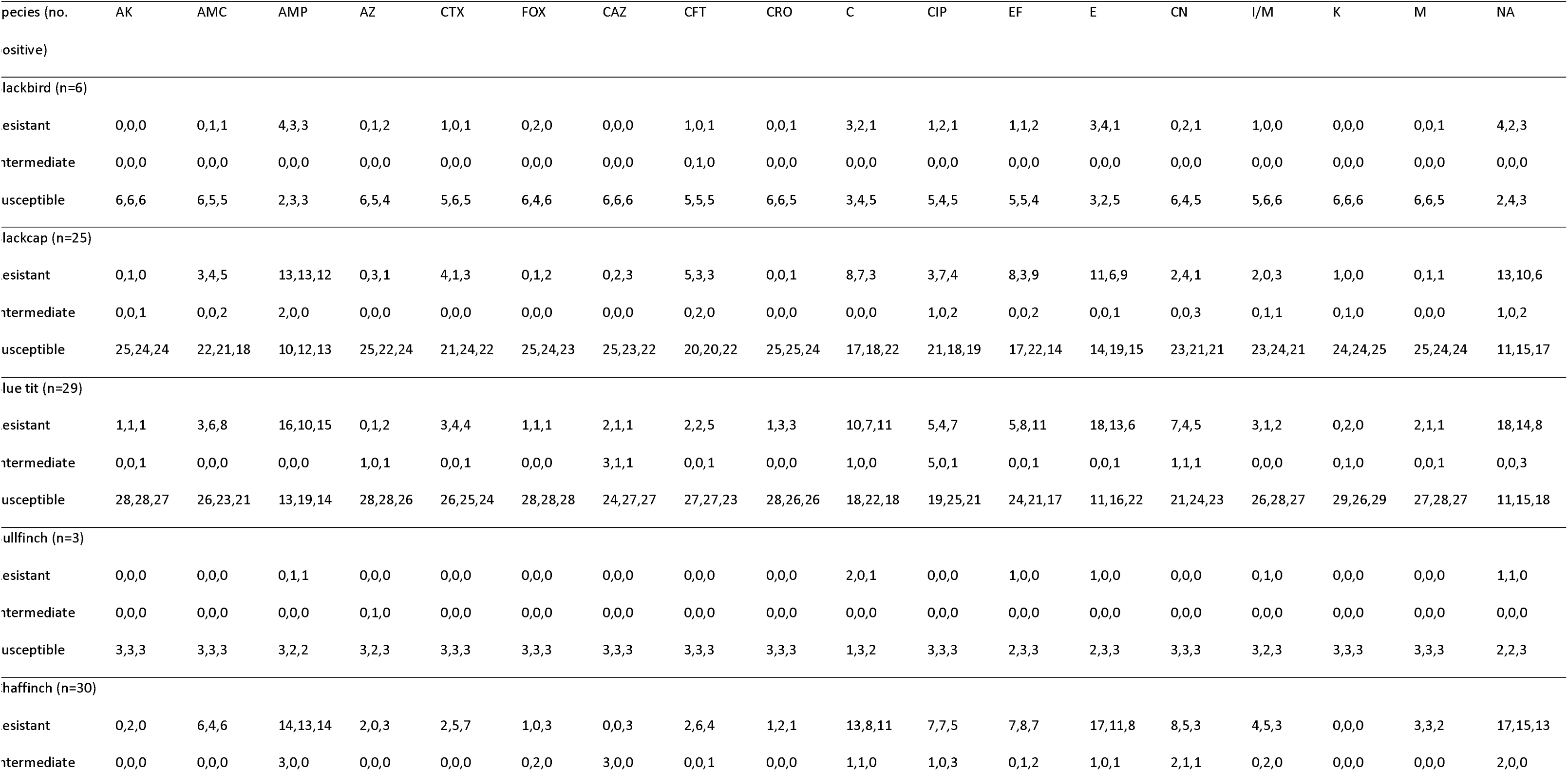
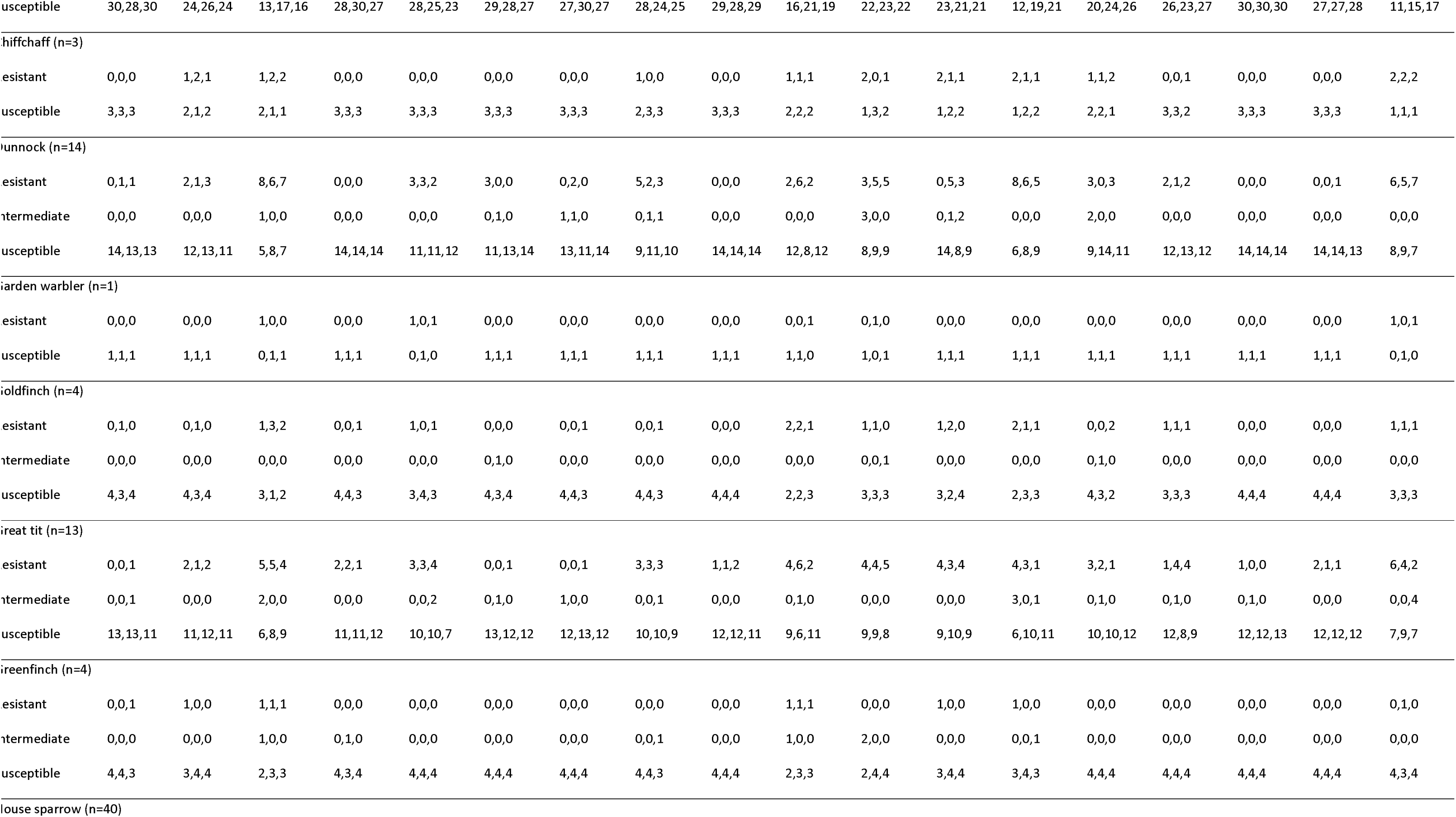
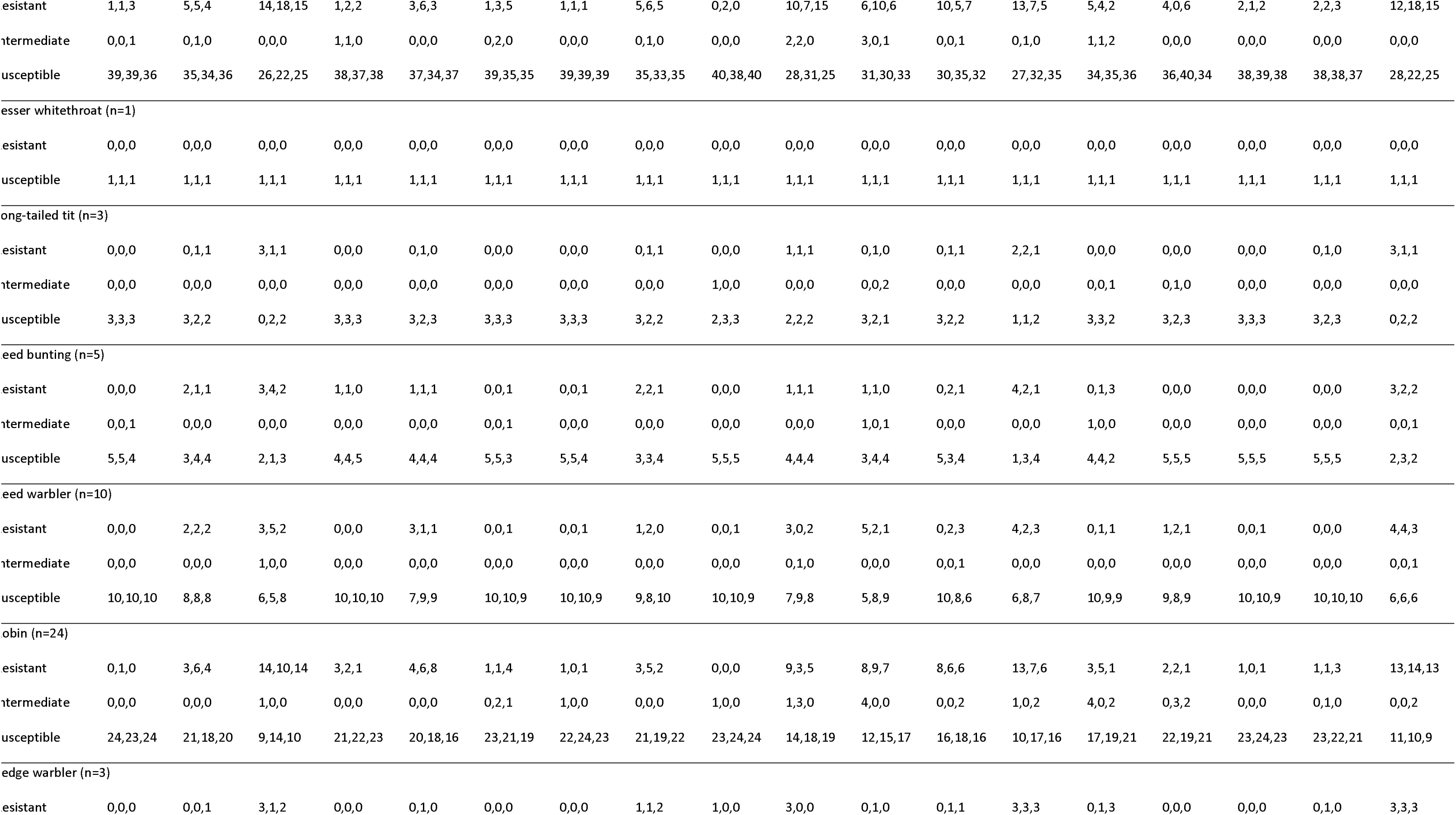
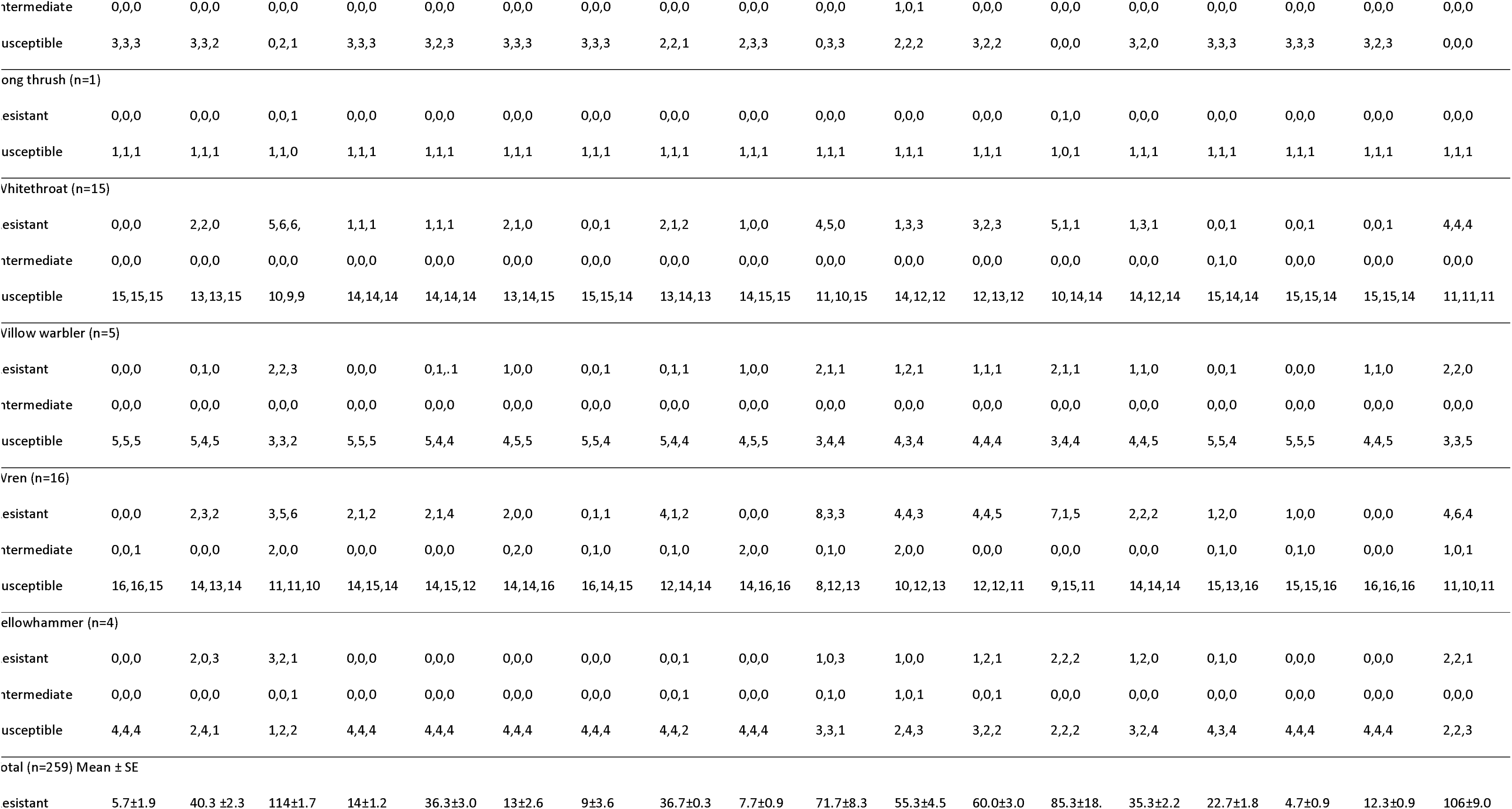
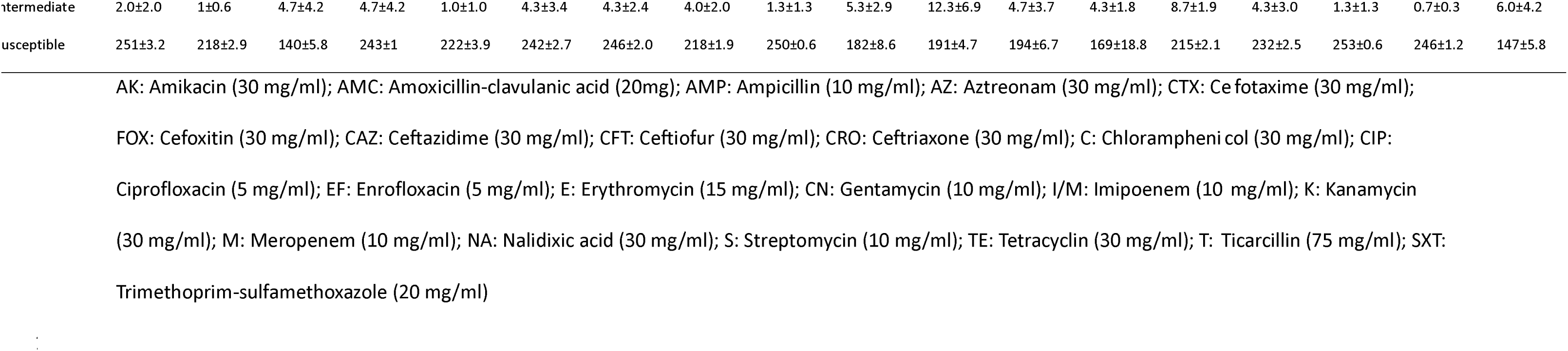
Antibiotic susceptibility of *E. coli* isolates recovered from passerine faecal samples. The three numbers in each cell represent three replicates; total data are the mean ± SE of the three replicate totals.

